# Global and local nature of cortical slow waves

**DOI:** 10.1101/2024.11.21.624645

**Authors:** J. Alegre-Cortés, M. Mattia, M. Sáez, R. Reig

**Affiliations:** Instituto de Neurociencias CSIC-UMH, Alicante, Spain; Natl. Center for Radiation Protection and Computational Physics, Istituto Superiore di Sanità, 00161 Rome, Italy

**Author notes:** Correspondence (J.AC), (R.R).

**Keywords:** Slow Wave Activity, computational model, sensitivity analysis, Up states, cortex

## Abstract

Explaining the macroscopic activity of a recorded neuronal population from its known microscopic properties still poses a great challenge, not just because of the many local agents that shape the output of a circuit, but due to the impact of long-range connections from other brain regions. Here we use a computational model to explore how local and global components of a network shape the Slow Wave Activity (SWA). We performed a sensitivity analysis of multiple cellular and synaptic features in models of isolated and connected networks. This allowed us to explore how the interaction of local properties and long-range connections shape the SWA of a population and its neighbors, as well as how the sequential propagation of active “Up” states lead to the emergence of preferred modes of propagation. We described relevant features of cortical Up states that are modulated by “stiff” combinations of parameters of the local circuit as opposed to other that are sensitive to the level of excitability of the whole network and the input coming from neighbor populations. We found that while manipulations in the synaptic excitatory/inhibitory balance can create local changes, cellular components that modulate the excitability or adaptation of a population have a long-range effect that leads to changes in neighbor populations too. Additionally, our simulations guided *in vivo* experiments that showed how heterogeneities in excitability between cortical areas can determine the directionality of travelling waves during SWA. We expect these results to motivate future research exploring and comparing cortical circuits through the analysis of their Up states.

## Introduction

One of the goals of neuroscience is to understand how circuits of neurons shape their activity. To achieve it, researchers have traditionally recorded neuronal activity during different behavioral states, coming from the early electroencephalogram recordings almost one century ago (Berger 1929) to our present toolbox that allows us to record the activity of up to many thousands of neurons simultaneously. Explaining how the recorded activity results from the interaction between the components of a circuit is not a solved question yet, considering that the output of a population is shaped not just by its own constraints, but because of the impact of long-range inputs from other brain areas. For example, two circuits may have identical properties but one of them has a higher firing rate if it receives a bigger amount of distant excitatory inputs.

We have aimed to improve our understanding of how spontaneous activity emerges from the combination of local and long-range components by means of manipulating cellular and synaptic parameters of *in silico* coupled neuronal populations under the slow wave activity (SWA) regime (Destexhe, Contreras, and Steriade 1999; Steriade, Nuñez, and Amzica 1993; Sanchez-Vives and McCormick 2000; Yousheng, Hasenstaub, and McCormick 2003; Sanchez-Vives, Massimini, and Mattia 2017), as a robust low-complexity brain state. This follows the idea that if the manipulation of a given component of a circuit can create discernable changes in the SWA, it may also be a major modulatory mechanism for the activity of that circuit across other behavioral states. SWA is generated during sleep and anesthesia, as well as abnormal conditions including isolated cortical slabs (Timofeev 2000) and perilesional areas (Russo et al. 2021). SWA can also be detected during quiescent awake periods (Mohajerani et al. 2010; Vyazovskiy et al. 2011; Nir et al. 2011). In this “default” activity pattern (Sanchez-Vives and McCormick 2000; Sanchez-Vives, Massimini, and Mattia 2017; Sanchez-Vives and Mattia 2014), the membrane potential of neurons alternates synchronously between hyperpolarized “Down” states and depolarized “Up” states of persistent high-firing activity, at a frequency of ∼1 Hz. Up states originate in L5 of the cortex (Sanchez-Vives and McCormick 2000; Compte et al. 2003; Beltramo et al. 2013; Wester and Contreras 2012; Capone et al. 2019), and from here propagate to other cortical and subcortical regions (Wilson and Kawaguchi 1996; Roš et al. 2009; Cowan and Wilson 1994; Timofeev and Steriade 1996; Sheroziya and Timofeev 2014; Stroh et al. 2013; Nir et al. 2011). The cellular and synaptic mechanisms that control the transition among these states have been widely studied *in vivo*, *in vitro* and *in* silico, as well as the possible roles of these oscillations in memory consolidation through their interaction with thalamic spindles (Steriade 2001; Nádasdy et al. 1999), or homeostasis (Tononi and Cirelli 2006; Vyazovskiy and Harris 2013). Multiple studies have also characterized in recent year how does the brain leave the SWA state during the process of wakening (Dasilva et al. 2021; Pazienti et al. 2022).

In particular, two components at the cellular and circuit level have been described as the main mechanisms controlling the emergence of Up states and their alternation with Down states during the SWA, the excitability of pyramidal neurons and their firing adaptation mediated by the accumulation of inhibitory currents during the Up state (Porta et al. 2023; Compte et al. 2003; Dalla Porta et al. 2024; Sanchez-Vives et al. 2010; Reig et al. 2010). Such microscopic mechanisms are amplified by the nonlinear dynamics of mesoscopic neuronal populations well-described by mean-field theories eventually giving rise to collective phenomena typically observed in relaxation oscillators (Latham et al. 2000; Gigante, Mattia, and Del Giudice 2007; Mattia and Sanchez-Vives 2012). In this framework the sleep-wake transition is determined by a weakening of the adaptation or complementary by a strengthening of the excitation receives by pyramidal neurons (Mattia and Sanchez-Vives 2012; Sanchez-Vives, Massimini, and Mattia 2017; Levenstein, Buzsáki, and Rinzel 2019; Jercog et al. 2017). Other studies have focused on a wider scale, exploring how long-range connections impact the propagation of the SWA and how the complexity of the cortical network can be explored studying the SWA, both in the untouched brain and through perturbations (Massimini et al. 2024; Idesis et al. 2024; Rosanova et al. 2018; Capone et al. 2019; Nghiem et al. 2020; Levenstein, Buzsáki, and Rinzel 2019; Mehrotra et al. 2023; Cakan et al. 2022). Despite all this progress in our understanding of the SWA, it remains an open question how the global network state and the local properties of a given population interplay to create particular dynamics during Up states. They are remarkably similar along the cortex during a traveling wave, yet some differences can be found across regions, i.e., in their tendency to initiate the waves, in the structure of the Up state or its firing rate (Ruiz-Mejias et al. 2011; Shimaoka, Song, and Knöpfel 2017; Greenberg et al. 2018; Pazienti et al. 2022). For example, somatosensory and motor areas have a higher tendency to present fractured Up states with a higher number of peaks, known as diplets (or triplets, quadruplets…) than other cortical areas (Alegre-Cortés et al. 2021). This Up structure is then preserved along other regions involved in somato-motor behaviors, as the dorsolateral striatum. Diplets can also emerge in other cortical regions upon bicuculline administration *ex vivo* slices (Sanchez-Vives et al. 2010).

How can we use Up state properties to unravel the differences among connected cortical circuits? To answer this question, we built a conductance-based model of coupled cortical L5 neuronal populations under the SWA regime, following and expanding the original work of (Compte et al. 2003). Conductance based models allow to consider the contribution of specific channels together with the widely described synaptic and network properties (Dalla Porta et al. 2023; 2024; Barbero-Castillo et al. 2021; Compte et al. 2003). We characterized which cellular and synaptic parameters can support locally particular Up states in a population that is part of a bigger network. We show how manipulations in the excitability and excitatory/inhibitory balance impacts the Up states, how this can be modulated by the presence of neighbor connections and propagate to them, including the different outputs that are driven by synaptic versus intrinsic inhibition. In addition, performing in vivo experiments we show how increasing the levels of cortical local excitability leads to changes in the preferred axis of propagation of slow waves in anesthetize mice Our sensitivity analysis of the SWA points at specific measurable parameters of the SWA that can be connected to mechanistically precise substrates, both at the local or the network level. According to predictions by mean-field theories (Mattia and Sanchez-Vives 2012; Jercog et al. 2017), we show how some parameters involving the excitability or adaptation of a population have a global effect, impacting their own SWA and their neighbor. On the other hand, certain levels of manipulation of both excitatory or inhibitory synaptic currents were able to create local changes in the Up states of the manipulated population only, impacting their length and number of peaks. This provides an improved understanding of how local and global components interact to modulate the activity of a neuronal population. We expect these results to provide a theoretical foundation for future studies; we discuss how our results can be extended to generate specific mechanistical predictions when different circuits are compared, as well as which specific activity motifs are good targets to study local properties, being relatively resilient to changes in the neighbor populations.

## Methods

### Ethical approval

All the experimental procedures were in conformity with the directive 2010/63/EU of the European Parliament and of the Council, and the RD 53/2013 Spanish regulation on the protection of animals use for scientific purposes, approved by the government of the Autonomous Community of Valencia, under the supervision of the Consejo Superior de Investigaciones Científicas and the Miguel Hernandez University Committee for Animal use in Laboratory.

### Electrophysiological recordings

#### *In vivo* extracellular recordings

Anesthesia was induced by an intraperitoneal injection of ketamine (75 mg/kg) and medetomidine (1 mg/kg). Briefly, after the induction, C57BL/6 mice (n= 9, 4 males and 5 females between 10 and 17 weeks old) were placed on a stereotaxic frame (Stoelting) with a tube blowing oxygen-enriched air placed ∼1 cm in front of their snout. The mice rested over a heating pad (FHC Inc.) to maintain body temperature at 36.5 ± 0.5°C. One third of the original induction dose was injected intramuscularly to maintain anesthesia once paw reflex could be evoked or the LFP trace exhibited signs of awakening, approximately every two hours. Extracellular activity was then recorded by two linear silicon probe electrodes (NeuroNexus Technologies, Ann Arbor, MI). One of them (A4×8-5mm-200-400-177) consisted of four shanks separated by 400 μm (8 electrodes in each shank spaced by 200 μm), which were inserted in a perpendicular angle of 38° to the cortex in the anterior region of the brain in the following coordinates: shank 1 (AP. +2.74mm); shank 2 (AP. +2.34mm); shank 3 (AP. +1.94mm); shank 4 (AP. +1.54mm). All of them with a LM. +2.7mm coordinate and a depth of 2.2mm. The other probe (A2×16-10mm-100-500-177) consisted of 2 shanks separated by 500 μm (16 electrodes in each shank spaced by 100 μm), which were inserted in a perpendicular angle of 38° in the primary visual cortex (V1) region of the brain in the following coordinates: shank 1 (LM. +3mm); shank 2 (LM. +3.5mm). Both with an AP. −3.75mm coordinate and a depth of 2mm. All coordinates followed (Paxinos and Franklin 2001).

Spontaneous activity was recorded before, during and after the extracellular local application of norepinephrine 50 μM (Abcam) and carbachol 5 μM (Tocris) diluted in NaCl 0,9%, by using a borosilicate micropipette, following (Barbero-Castillo et al. 2021; D’Andola et al. 2018). Its tip was introduced in the same craniotomy, next to the probe located in the V1 region (n=9). In 5 animals, after delivery of norepinephrine and carbachol and once activity returned to normality, NaCl 0,9% was delivered in similar fashion for control purposes. Data was continuously digitized at 20 kHz using a SmartBoxV2 amplifier (NeuroNexus Technologies, Ann Arbor, MI).

After each experiment, probes were stained with 1,1′-Dioctadecyl-3,3,3′,3′-tetramethylindocarbocyanine perchlorate (Dil, Sigma) for at least ∼15 minutes and then lowered to the previous position. There, they were maintained for at least ∼20 minutes. Afterwards, animals were sacrificed with a lethal dose of sodium pentobarbital (150 mg/Kg) and perfused with a solution containing 4% paraformaldehyde in 0.1 M phosphate buffer (PB, pH 7.4). Brains were extracted and stored in PBS solution until the cutting. Before processing, brains were transferred into PBS containing 30% sucrose for at least 24/48 hours. Brains were then cut with a criotome (Microm) in 50 μm sections and collected on gelatin coated slides (ThermoFisher), covered with mowiol (Calbiochem) and then mounted with coverslips (ThermoFisher). Finally, we imaged them using a fluorescence wide field microscope (Thunder, Leica).

### Electrophysiological data analysis

#### Detection of Up and Down states

We used the 4 shanks placed covering rostro-central cortical areas (FrA, M1 and S1) and an extra shank from the probe in V1 that lied in the same rostrocaudal plane. We selected the electrodes placed in the cortex based on their distribution along the probe and insertion depth. Then we divided the recorded local filed potentials (LFP) into Up and Down states after estimating the multiunit activity (MUA) (Reig et al. 2010; Mattia, Ferraina, and Del Giudice 2010). To do so, we first computed the MUA as the power of the signal in the 200-1500 Hz band using 5 ms windows. We then computed the logMUA as the log(MUA) and smoothed it using 80ms bin. We then separated Up and Down states using a threshold calculated as the mean +0.5 std of the signal. When two Up states were detected with less than 250 ms of separation they were merged in a unique Up state. At last, transitions with durations shorter than 100 ms were removed from the Up states pool.

#### Slow Waves detection and reconstruction

We computed global Up states following the analysis of (Capone et al. 2019). In brief, we used a rolling window with an initial length of 1s, which was shrined iteratively by a factor of 0.75 until if contained no more than 1 transition from Down to Up states per electrode. We then selected those windows on which an Up state was detected in at least electrode of each shank for further analysis, considering them global Up states. For each global Up state, we computed the delay between the transition detected in each electrode and its average across electrodes. These values were stored in a time lag matrix of size [Electrodes x Global Ups] which was store for further analysis.

#### Network model

Each population in our model is composed of 1024 excitatory and 256 inhibitory neurons randomly distributed along a line and connected through biologically plausible synapses. The model mostly follows (Compte et al. 2003), with some parameters borrowed from (X. J. Wang and Buzsáki 1996; Barbero-Castillo et al. 2021). Any slight modification to the original conductance values is detailed in Table 1.

**Table 1.**
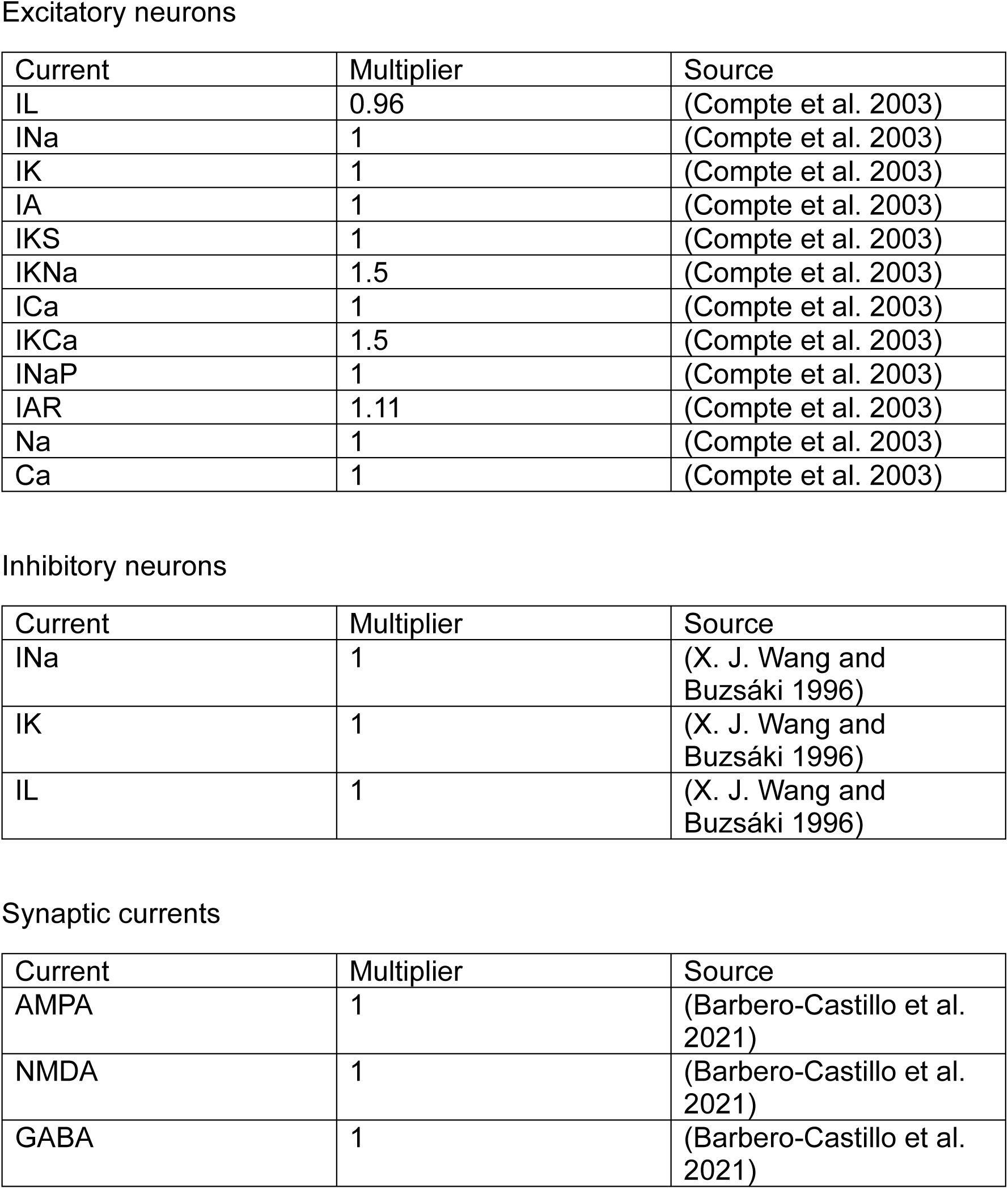
Source of the parameters used in the model and their modifications.

Excitatory neurons
| Current | Multiplier | Source |
| --- | --- | --- |
| IL | 0.96 | (Compte et al. 2003) |
| INa | 1 | (Compte et al. 2003) |
| IK | 1 | (Compte et al. 2003) |
| IA | 1 | (Compte et al. 2003) |
| IKS | 1 | (Compte et al. 2003) |
| IKNa | 1.5 | (Compte et al. 2003) |
| ICa | 1 | (Compte et al. 2003) |
| IKCa | 1.5 | (Compte et al. 2003) |
| INaP | 1 | (Compte et al. 2003) |
| IAR | 1.11 | (Compte et al. 2003) |
| Na | 1 | (Compte et al. 2003) |
| Ca | 1 | (Compte et al. 2003) |

Inhibitory neurons
| Current | Multiplier | Source |
| --- | --- | --- |
| INa | 1 | (X. J. Wang and Buzsáki 1996) |
| IK | 1 | (X. J. Wang and Buzsáki 1996) |
| IL | 1 | (X. J. Wang and Buzsáki 1996) |

| Current | Multiplier | Source |
| --- | --- | --- |
| AMPA | 1 | (Barbero-Castillo et al. 2021) |
| NMDA | 1 | (Barbero-Castillo et al. 2021) |
| GABA | 1 | (Barbero-Castillo et al. 2021) |

Neurons are homogeneously distributed along a line of 5 mm and sparsely connected to each other using a Gaussian probability function centered at each neuron and following the same distribution as in (Compte et al. 2003). Each neuron in a given population contacted an average of 95+/-12 neighbors. Connected populations were coupled thorough long-range excitatory connections with a probability of 0.008, which represents a distance between populations of ∼500 µm (Potjans and Diesmann 2014) and targeting both excitatory and inhibitory neurons (Haider and McCormick 2009; Mcguire et al. 1991).

#### Model implementation and simulation

The model was implemented using Brian 2 (Stimberg, Brette, and Goodman 2019) in Python and simulated using a fourth order Runge-Kutta method with and adaptative time step.

#### Simulations analysis

To study the subthreshold activity of the simulated populations, 100 traces from randomly selected excitatory neurons were downsampled to 1 KHz and stored for analysis. Then Up and Down states were detected by first smoothing the signal using a 100 ms window and then applying a threshold computed as the 60% distance between the peaks of two gaussian functions fitting the bimodal distribution of the histogram of Vm values (Sanchez-Vives et al. 2010). If the detection of the two peaks in the distribution failed, we considered that neuron to now follow a bimodal distribution and excluded it from further analysis of the Up states. When two Up states were detected with less than 250 ms difference, they were merged into a single Up state. Up states shorter than 250 ms were excluded. We then computed the amplitude, length, slope speed from the Down to the Up states as well as Up to Down states, number of peaks and peak amplitude of the Up states following (Alegre-Cortés et al. 2021). Slopes were computed using a 100 ms (Down to Up) or 60 ms (Up to Down) time window around the state transitions, we computed the derivative of the time series and smoothed it using a 10 ms window. We then computed the absolute value of the derivative and delimited the transition using a threshold consisting of the mean + 1 std. Once the transition between states was found, a liner regression was fitted to compute the slope speed. Peaks were computed using the *findpeaks* function from *Scipy* package in the Up state, with a peak height of 0.3 std and a minimum distance between peaks of 160 ms. Peak distance was measured as the distance in mV between the minimum and maximum value of the Up state. qSpiking activity was directly stored as time stamps as an output of the simulations and analyzed without further pre-processing.

## Results

### Strategy and computational model

In order to understand to what differences in the multiscale properties of a defined neuronal population modulated both its output and the one of its neighbors we sequentially manipulated multiple intrinsic and synaptic properties of isolated and coupled populations *in silico*.

Cortical recordings show a homogeneous frequency of appearance of Up states as well as a preferred rostro-caudal directionality in mammals under deep sleep/anesthesia (Massimini et al. 2004; Pazienti et al. 2022; Alegre-Cortés et al. 2021). We predicted that long-range connections could propagate Up states from highly excitable areas, or “hot spots”, blurring a potentially heterogeneous landscape of local excitability levels. In order to be able to test this hypothesis in silico, we decided to manipulate the leakage current which, among other mechanisms, contributes to control neuronal excitability due to of its effect on the voltage during the Down state. (Fig. 1g,h). This inward rectification current is active during the Down state and exerts a control of the membrane voltage during, therefore influencing the required force to move the neuron into the active Up state. We characterized how the frequency of oscillation is modulated by means of the leak current (Fig. 1h) and manipulating the conductance of this channel we combined populations with different frequencies of oscillations in the following sections.

**Fig. 1.**
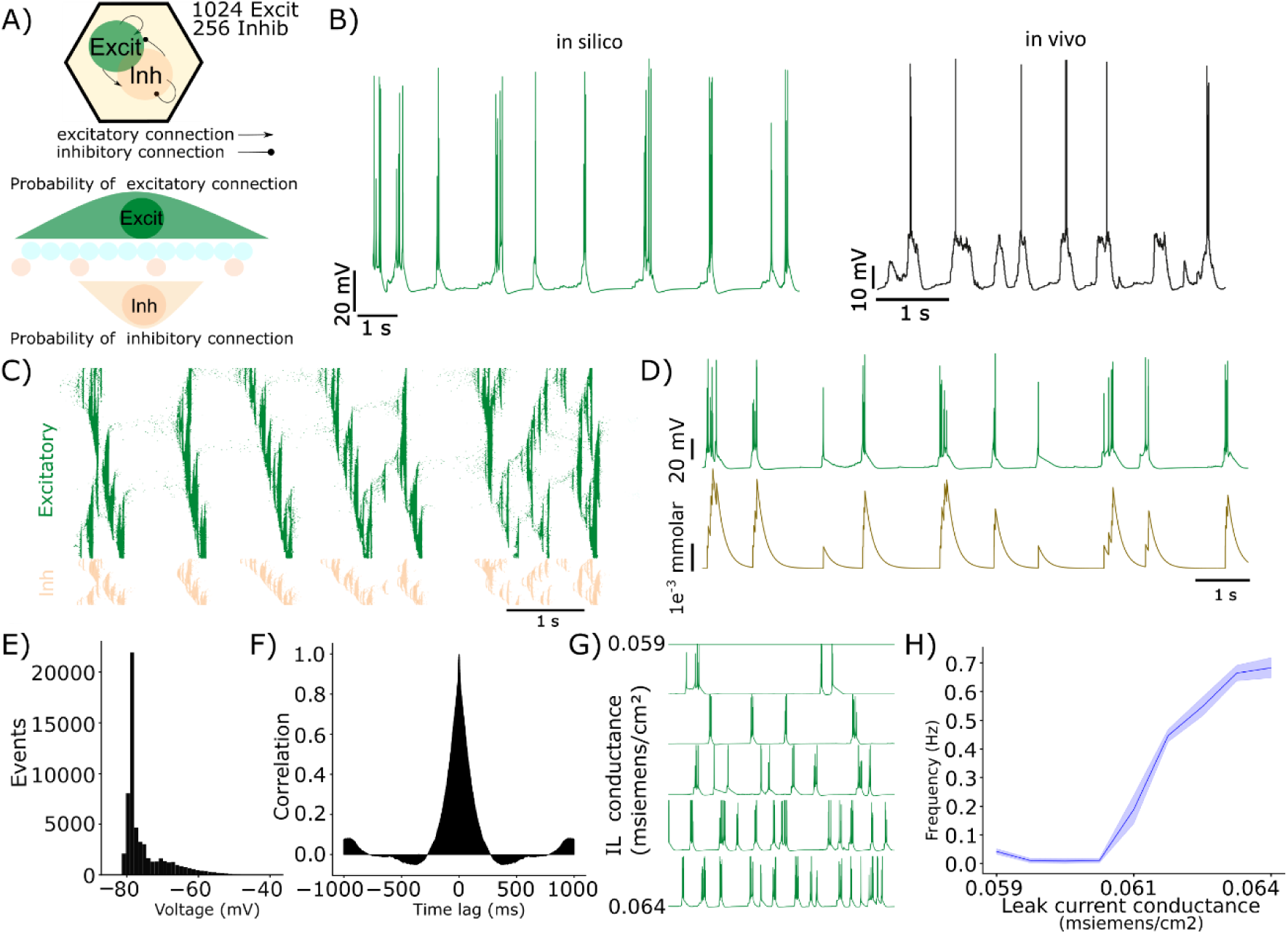
SWA computational model and excitability. **A)** Schematic representation of the components of the model showing the cell types and probability distribution of synaptic contacts. **B)** Representative examples of the spontaneous activity under the SWA regime of an excitatory neuron of the model (left) and Layer V pyramidal neuron recorded in whole-cell patch-clamp (right). **C)** Raster plot visualization of a modelled population, each row represents a neuron and each point a spike. Spikes from excitatory neurons are shown in green and from inhibitory neurons in light orange. **D)** Spontaneous activity of an excitatory neuron (top) of the model and its respective intracellular calcium concentration (bottom). **E)** Histogram of membrane voltage of an excitatory neuron. **F)** Autocorrelogram of the membrane voltage of an excitatory neuron. **G**) Impact of the conductance of the leak current in the frequency of the SWA. **H)** Relationship between the conductance of the leak current and the average frequency of the population up states.

### Propagation of Up states and homogenization of oscillatory frequency

During deep sleep or anesthesia, SWA appears as travelling waves along the cortex and from here to other brain regions. During the travelling wave, populations are sequentially activated and thus show a similar frequency of Up states. We first asked whether this state comes from a similar level of excitability along all the cortical populations, or if Up states can propagate from “hot spots” and evoke new Up states in populations that otherwise would lack the excitability to transition to Up states autonomously or would do in a longer timescale.

We connected two populations with a different intrinsic frequency of oscillation simulating a distance of approximately 500 μm (Fig. 2a), following previous works that estimate the probability of pyramidal neurons stablishing synaptic contacts onto other neurons (excitatory and inhibitory) based on their distance. The model does not include long-range projections from inhibitory neurons. When a raster plot is visualized to compare the firing dynamics of the two pairs of populations (Fig. 2b, connected vs disconnected), the slow population increases its frequency of Up states to match the one of the populations with a higher frequency. As previously described in vivo (Ruiz-Mejias et al. 2011), a small group of neurons could be seen preceding the Up state in the leading population. We then quantified the frequency of oscillation of the two populations, comparing them connected and disconnected, and gradually decreasing the excitability of the slow population. Dashed lines in Fig. 2c represent the frequency of oscillation of the two populations when they are disconnected. The frequency of the slow population (dashed green) increases with the conductance of the leak current up to a point on which it is the same as the high frequency population (dashed purple). This later one remains untouched during the simulations, thus why its frequency remains unchanged. On the other hand, when both populations are coupled, the changes in the conductance of the leak current of the slow population do not have an impact in the oscillatory frequency anymore, aligning to the frequency of oscillation of the most excitable one. Notably, connecting the populations did not clearly increase the overall frequency of up states. This suggests that the frequency of the SWA of a network may be modulated by the most excitable population.

**Fig. 2.**
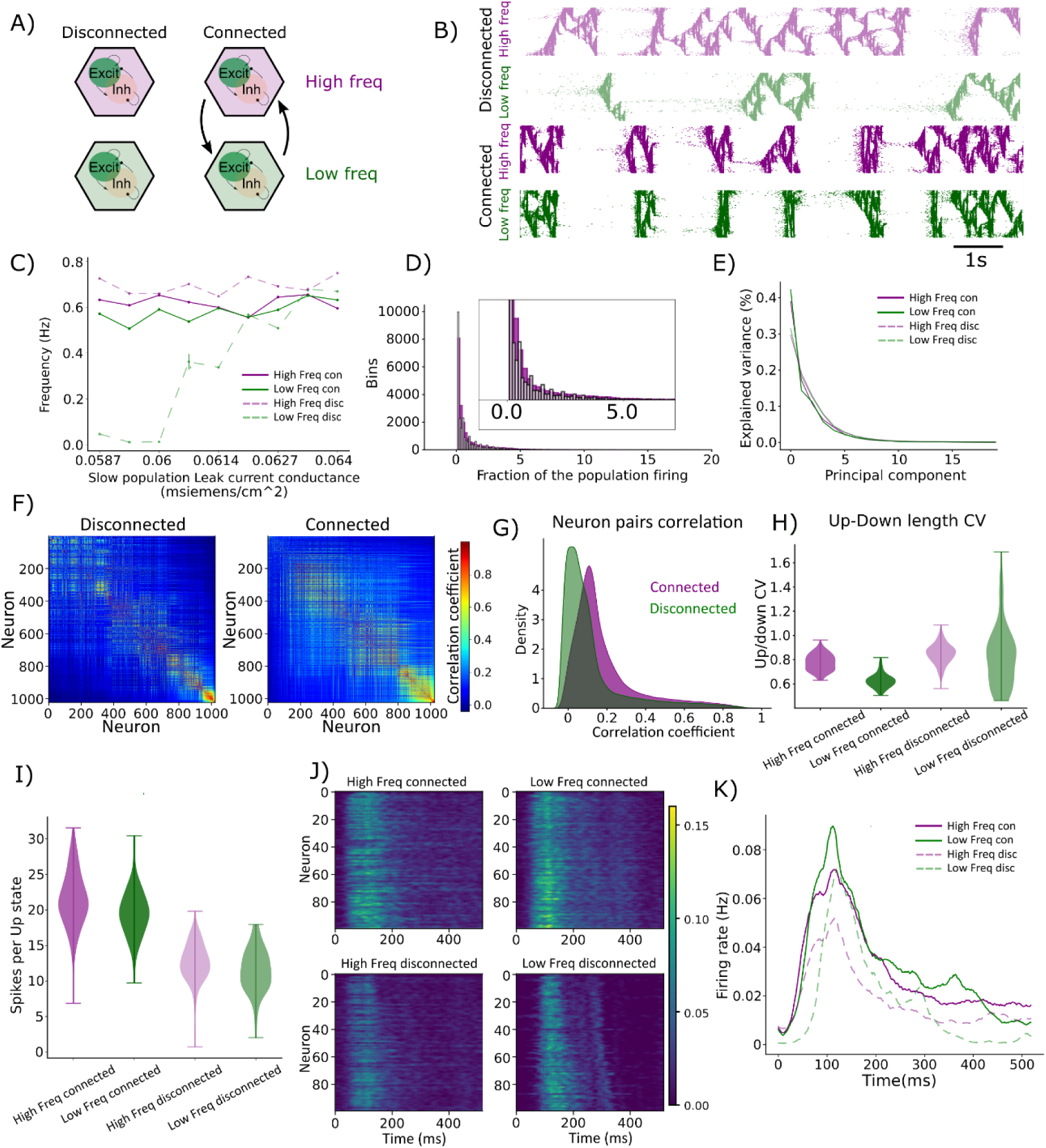
Impact of long-range connections in a model of SWA. **A)** Schematic representation of the different simulations. Each of them includes an unperturbed population that oscillates at a fixed higher frequency and a second one with a decreasing frequency of oscillation across simulations. Both populations are simulated with and without long-range connections between them. **B)** Raster plot of both populations in the absence (top) or presence (bottom) of long-range connections. When connections are present, they synchronize the transition to the Up states following the rhythm of the one with higher frequency. **C)** Frequency of the SWA of the higher and lower frequency populations before and after long-range connections are included in the model. The slower population oscillates at the rhythm of the faster one when they are coupled, independent of its original frequency. On the other hand, the population of higher frequency is not affected by the addition of the long-range connections. **D)** Histogram showing the fraction of neurons firing together (5 ms window) in a connected vs disconnected population. The presence of long-range connections increases the number of neurons firing together and therefore the synchronicity of the population. **E)** Percentage of variance explained by each PC when a PCA is computed on the firing activity of a connected and a disconnected population. Long-range connection increase the variance explained by the 1^st^ PC. **F)** Correlation matrix of a disconnected and a connected population. **G)** Connected populations (purple) have a higher average correlation coefficient between pairs of neurons than disconnected. **H)** Rhythmicity is increased by long-range connections. Long-range populations show a reduced Up/down coefficient of variance. **I)** Long-range connections increase the firing rate of the populations. **J)** Raster plots of each type of simulated population. Each row shows the average firing rate of one neuron, aligned to the onset of the Up state. **K)** PSTH of the data shown in J). The increase in firing on the connected populations is due to an overall increase in the number of spikes along the Up state.

We then explored the impact in the firing rate of the populations caused by the introduction of the long-range connections between them. First, we quantified this as the number of neurons firing simultaneously in bins of 5 ms, when the population was connected or disconnected from the other (Fig. 2d). A shift in the histogram is visible, with a higher fraction of neurons firing simultaneously when populations are connected. We then explored if this increase in synchronization led to a change in the dimensionality of the population dynamics by means of computing the weight of the first principal components of the spiking activity of all the neurons in the four populations (Fig. 2e). The first component of the connected populations explains around 40% of the total variance, while a 30% of the disconnected populations. This implies a relative increase of ∼33% of the amount of variance explained by the first PC, in line with previous studies using recurrent neuronal networks, on which cross-region communication quenched local dimensionality (Clark and Beiran 2024). Consistently, neurons in the connected populations were higher correlated among them than in the disconnected ones (Fig. 2f,g).

The reduction in the dimensionality of the SWA when two populations were connected occurs together with an increase in the rhythmicity (Fig. 2h), computed as a decrease in the coefficient of variance of the length of each full cycle. This effect was particularly notorious in the case of the slower populations (0.82±0.25 vs 0.63±0.06).

Lastly, we explored how connecting the populations impacted the firing rate. Once again, the main difference of firing was between connected and disconnected populations (Fig. 2i, 21.4±4.1 Hz and 19.7±3.6 Hz vs 12.6±2.9 Hz and 11.5±3.0 Hz), while the differences due to changes in excitability were marginal. Considering that the populations were more synchronized, we asked whether the increment in firing rate was due to a bigger peak of firing rate, a small window with a higher number of spikes in the population, or due to an increment in the firing rate along the Up state. To do so, we first computed the Up and Down states of the first principal component of the population firing (see methods) and then the average PSTH of each neuron aligned to the transition to the Up states in the first PC (Fig. 2j). We then averaged the PSTHs to have a measure of the distribution of spikes during the Up states of each population (Fig. 2k). It shows how the increase of firing rate that follows the introduction of long-range projections in the model is caused by an increase in the number of spikes along the whole Up state. Note how the follower population has a sharper transition to the Up state together with a higher initial peak, because of receiving an Up state in opposition to creating it.

In summary, we explored the impact of coupling cortical populations under the SWA regime. We showed how differences in local excitability are blurred when populations were connected. The population with the higher excitability will impose its own rhythm hindering to decipher the differences in excitability from the firing rate of the populations once they were connected.

### Excitability hotspots govern SWA propagation axis in vivo

We have shown *in silico* that directionality was caused by differences in the levels of excitability between populations, implemented as changes in the conductance of the leak current. To test this prediction in vivo, we performed multielectrode recordings in multiple brain areas, while locally increasing the excitability of the caudal cortex by releasing a cocktail of norepinephrine (NE) and carbachol (CCh) diluted in sodium chloride (0,9%). Previous in vitro studies (D’Andola et al. 2018; Barbero-Castillo et al. 2021) have shown how SWA accelerated by adding these compounds.

A schematic representation of the experimental setup can be seen in Fig. 3a; we combined two multisite electrodes to cover the frontal and caudal regions of the cortex. A first neural multisite probe (Neuronexus) consisting of 4 shanks with 8 channels distributed in a column each was inserted parallel to the midline, while a second probe with 2 or 4 shanks was inserted in visual cortex. We recorded SWA activity from anesthetized mice (Fig. 3b) and isolated Up and Down state based on the logarithm of the multiunit activity (logMUA, Fig. 3c) from the electrodes placed in the cortex. During the recording, Ne and CCh were released in the proximity of the caudal recording site using a micropipette connected to a puffer (see methods). The release induced a change in the directionality of the SWA, which was initially rostrocaudal (Fig. 3d,e). To quantify directionality, we chose the caudal electrode that was placed in the same rostrocaudal plane as the rostral shank. When we isolated each wave (see methods) and computed the mean delay of each electrode, we could see how the delay was initially positive (following the wave) and shifted to negative values following the release (leading) during several minutes (Fig. 3d). When a matrix representing the relative onset delay of each Up state recorded in at least one cortical channel of each shank the transition from rostrocaudal to caudorostral predominant axis of propagation becomes clear in Fig. 3e. In addition, the release of NE and CCh induced an increase in the frequency of the SWA in the caudal cortex, which, consistently with the predictions of our model, lead to an increase of the frequency of the SWA in all the recorded regions (n=9, 0.60±0.26 vs 0.73±0.17 Hz, p value = 0.039) A representative example is shown in Fig. 3f. When we compared the fraction of rostrocaudal or caudorostral Up states before and after release (n=9), we saw a significant decrease of rostrocaudal Up states (n=9, 0.67±0.15 vs 0.5±0.22 p value = 0.0391), which was not visible in control animals (n=5, 0.69±0.14 vs 0.63±0.18, p value = 0.31). Based on this result, we validated the model prediction that a population of higher excitability leads cortical waves and imposes a common frequency of Up states into less excitable areas.

**Fig. 3.**
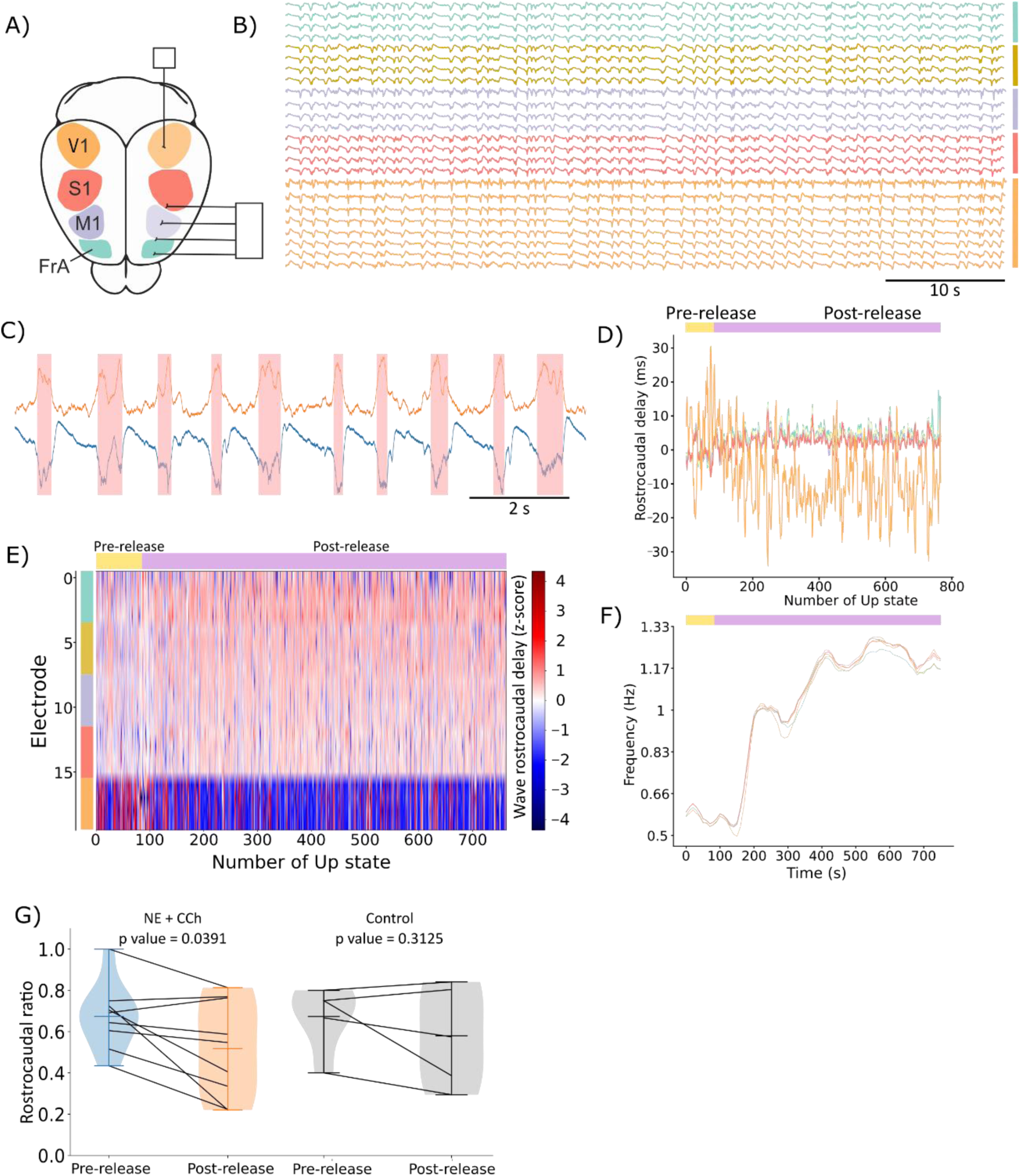
SWA propagation axis are controlled by excitable populations *in vivo*. **A)** Schematic representation of the *in vivo* recording setup. **B)** Representative example of the SWA activity recorded. Each color represents and individual shank, placed as shown in A). Shaded traces were recorded in the cortex and used for further analysis. **C)** Example of the detection of Up states, shown as faint red bars, based on the calculation of the log MUA of the recorded trace, shown in orange and blue respectively. **D)** Relative delay of each shank along the Up states of one recordings session. Negative values are shown in the shanks on which the Up state was recorded first. Note how after NE+Cch are released, the more caudal shank (orange) shifts from positive to negative delays, while the contrary occurs for the rest of the shanks, as the wave becomes predominantly caudorostral. **E)** Time lag matrix showing all the cortical electrodes and Up states recorded in a session. Note how following the release the caudal shank (orange) shifts from positive to negative delays, meaning that it becomes the first shank on which the Ups are detected. **F)** Following the release, the frequency of the SWA is increased in all the recorded shanks. Representative example. **G)** Change in the ratio of rostrocauldal Up states following the release in all the recorded sessions. There is a significant (p = 0.039) decrease in the amount of rostrocaudal Up states. Control recordings, on which saline was released as a substitute of the NE+CCh cocktail, do not show a significant change in the directionality of the SWA.

### Impact of connecting populations on Up state parameters

We next explored the impact of connecting populations on their subthreshold activity. Conductance based models display complex membrane voltage dynamics based on the interaction of the different channels, allowing to explore their role in different phases of the SWA. As shown at the level of populational spiking activity, the membrane voltage of the neurons of the fast and slow population became aligned when long-range projections were added to the model (Fig. 4a).

**Fig. 4.**
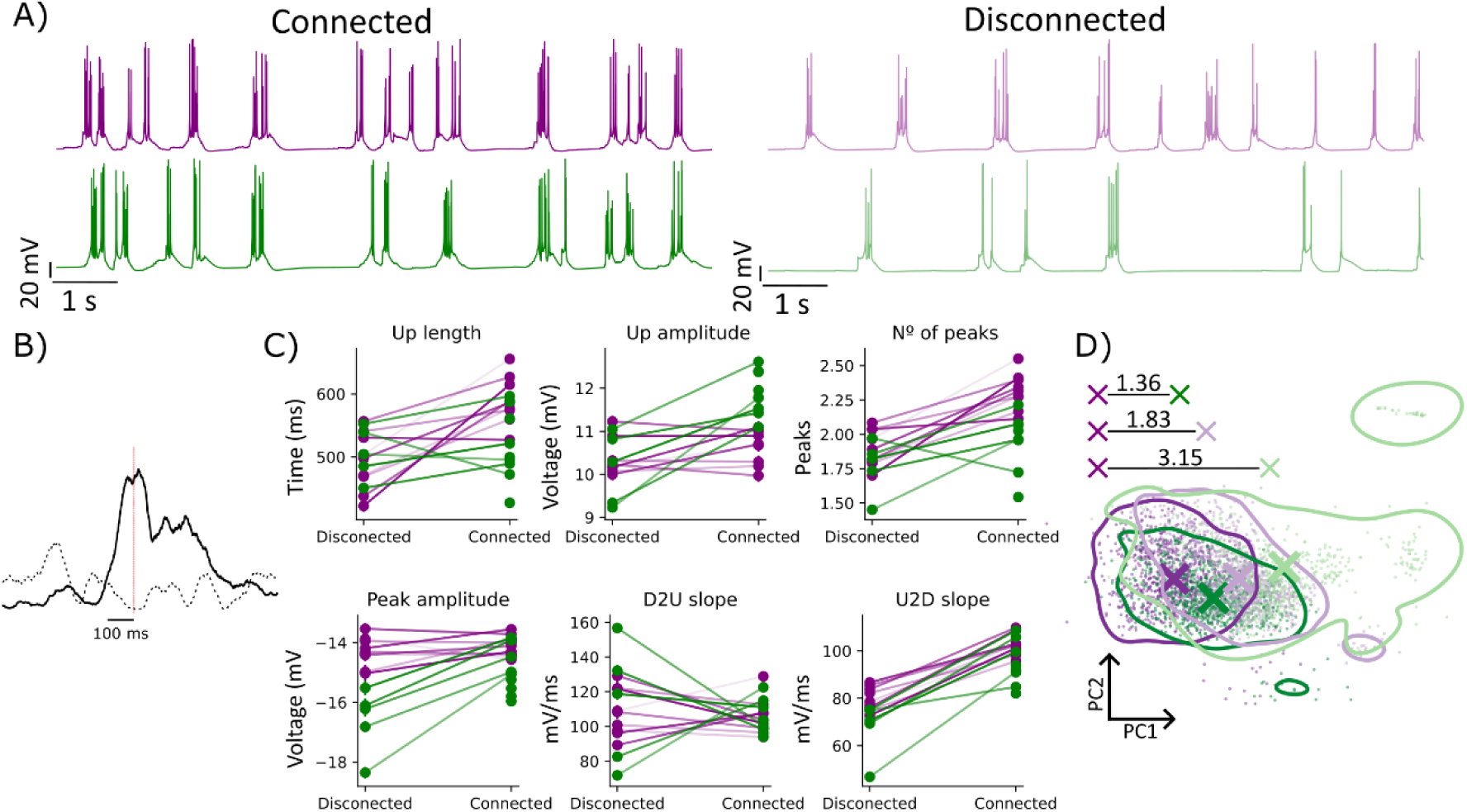
Impact of long-range connection on subthreshold dynamics. **A)** Examples of the membrane voltage of two neurons from two populations with higher (purple) and lower (green) frequency of oscillation when they are connected (left) or disconnected (right). **B)** Histogram of the transitions to the Up state of the slow population aligned to the transitions to the Up state of the faster population in the absence (dashed black line) or presence (black line) of long-range connections. **C)** Impact of long-range connections in the properties of the Up state. Simulations were performed with and without long-range connections and using decreasing values of conductance of the slow population (green) while keeping the faster population unmodified (purple). The level of fainting of the connecting line is proportional to the decrease in the conductance of the slower population in that simulation. **D)** Representation in the space conformed by the 2 first PC after performing PCA to the parameter space (shown in C) computed from all the excitatory neurons of all the simulations.

We computed the transition to the Up states of a random subsample of 100 neurons aligning the Up states of the fast and slow populations, from simulations with and without long-range connections (Fig. 4b). When the distribution of transitions to the Up state of the slow population is aligned to the faster population, most of the distribution falls to the positive side, confirming that Up states propagate from the fast to the slow population. We then quantified the impact of long-range connections on the Up states (Fig. 4c). As in Fig. 2, we simulated two populations with different excitability by gradually decreasing the leak current conductance of one of them while keeping the properties of the other one intact. Connecting the populations had an effect generally associated with an increase in excitability, with an elongation of the Up state and higher speed of the transitions from Down to Up and Up to Down states of both populations (Fig. 4c). The amplitude of the Up states and peak amplitude (see Methods) of the Up states of the slower population, as well as the number of peaks of the Up states of both populations. Interestingly, connected populations displayed similar values, with the only visible difference of an increment of >2 mV in the Up states’ amplitude of the slow population. In line with our previous results (Fig. 2) this demonstrates how long-range connections tend to homogenize the shape of the Up states, hindering the study of their local properties. To further explore this, we projected the neurons of the four populations and all simulations into the subspace of the two principal components, after applying a PCA to the parameters space. Fig. 4d shows the distance between the centroids of each population, using the fast connected population as reference. The most similar population (closest centroid) was the slow connected population. Note how we combined all the simulated traces, thus the slow populations (connected and disconnected), include simulations with a decreasing level of leak conductance. Despite this, connected populations were more similar among them than connected a disconnected population with the same conductance of the leak current. This confirms that the presence of long-range connections imposes a certain type of Up state into the network, determined by the most excitable population.

### Sensitivity analysis

We have shown a tendency to homogenize SWA after connecting neuronal populations. Nevertheless, previous works demonstrated local differences between cortical areas. For example, the internal structure of the Up states, in somato-motor regions displays diplets, defined by sharp deflections of the membrane voltage (Alegre-Cortés et al. 2021) or in the spontaneous action potentials, with higher frequency in prefrontal cortex (Ruiz-Mejias et al. 2011). Given the robustness and low-dimensionality of the SWA (Sanchez-Vives, Massimini, and Mattia 2017), we propose that those cellular and synaptic mechanisms that are capable of produce perceptible differences in the local properties of Up states are potential key regulators of the circuit.

We run different simulations manipulating the conductance of outward rectification, glutamatergic and GABAergic currents in a range spanning from 50% to 150% of their original value (Figs. 5-8). Each simulation consisted of three populations. One population was manipulated and had no long range connections, while a second population was manipulated in an identical fashion but was coupled through long-range connections to a neighbor, unmanipulated, population. This configuration followed two porpoises: First, we could explore the influence of long-range connections in the result of the manipulations. Second, we could see whether the manipulations impacted only the targeted population or had also a discernable impact on the neighbor, in the case of the coupled populations. We considered as local those changes that were visible only in the manipulated connected population and global if they were also present in the neighbor.

**Fig. 5.**
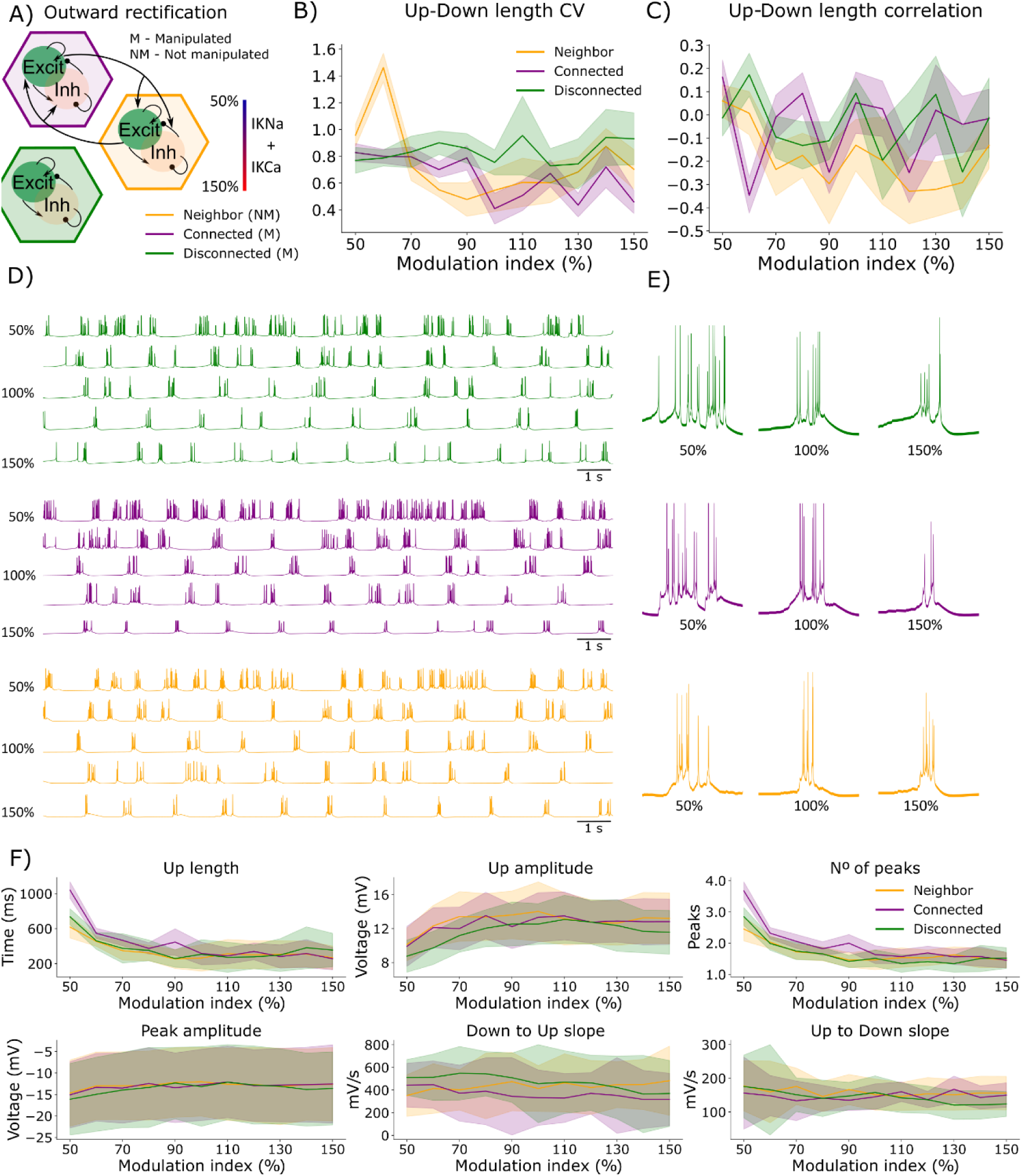
Impact of the manipulation of Ca^2+^- and Na^+^-activated K^+^ current conductance on SWA. **A)** Schematic representation of the simulations **B)** Coefficient of variance of the cycle length for the different levels of manipulation. **C)** Correlation of the length of each Up state and the following Down state for the different levels of manipulation. **D)** Representative traces of the neurons in the three manipulations shown in A) during different levels of manipulation (from 50% to 150%). **E)** Up states examples of the traces in D). Each trace is 1 second long. **F)** Impact on the properties of the Up states resulted from the simulations shown in A).

### Outward rectification

To begin, we manipulated intrinsic rectification, Na^+^ and Ca^2+^ activated K^+^ currents (Fig. 5), given their relevance in the control of the Up state (Compte et al. 2003). This manipulation led to an increase in the cycle CV of the connected populations when the conductance of these currents was close to 50% (Fig. 5b), with a particularly strong input in the neighbor unmanipulated population, and no change in the disconnected one. While there was not an impact in the relationship between the duration of each state within a cycle (Fig. 5c), the manipulation of Na^+^ and Ca^2+^ activated K^+^ currents had a perceptible impact on the Up states (Fig. 5d-f).

Increasing the conductance of these currents in the explored range, up to 50%, had a neglectable effect on the Up state properties, except for a decrease in the speed of both slopes in the disconnected population. On the other hand, decrease their conductance had a strong impact on the manipulated populations as well as the connected neighbor. All the populations showed an elongation and a decrease in the number of peaks of the Up states, and a decrease in their amplitude that was related to the decrease in the conductance. The two manipulated population additionally showed an increase in the speed of the Down to Up transition that was not present in the neighbor population. These results indicate that the manipulation of the outward rectification channels leads to global changes that propagate to the neighbor populations. Consistent with the strong global effect, the reduction of the conductance of the outward rectifiers shifted the dynamic state of all the populations to a partially asynchronous regime with long lasting depolarizations (Fig. 5d) that resembles the state before wakefulness.

### Glutamatergic currents

The explored range of manipulation of the conductance of AMPA and NMDA synapses together (Fig. 6a) was directly connected to changes in the SWA regime. The coefficient of variance in the length of each state within a cycle (Fig. 6b) was not clearly modulated by the manipulation. Nevertheless, decreasing the glutamatergic driving force led to a SWA of increased rhythmicity, with a lower correlation of Up and Down lengths (Fig. 6c). Representative traces and Up states examples along the range of manipulation are shown in Fig. 6d and e.

**Fig. 6.**
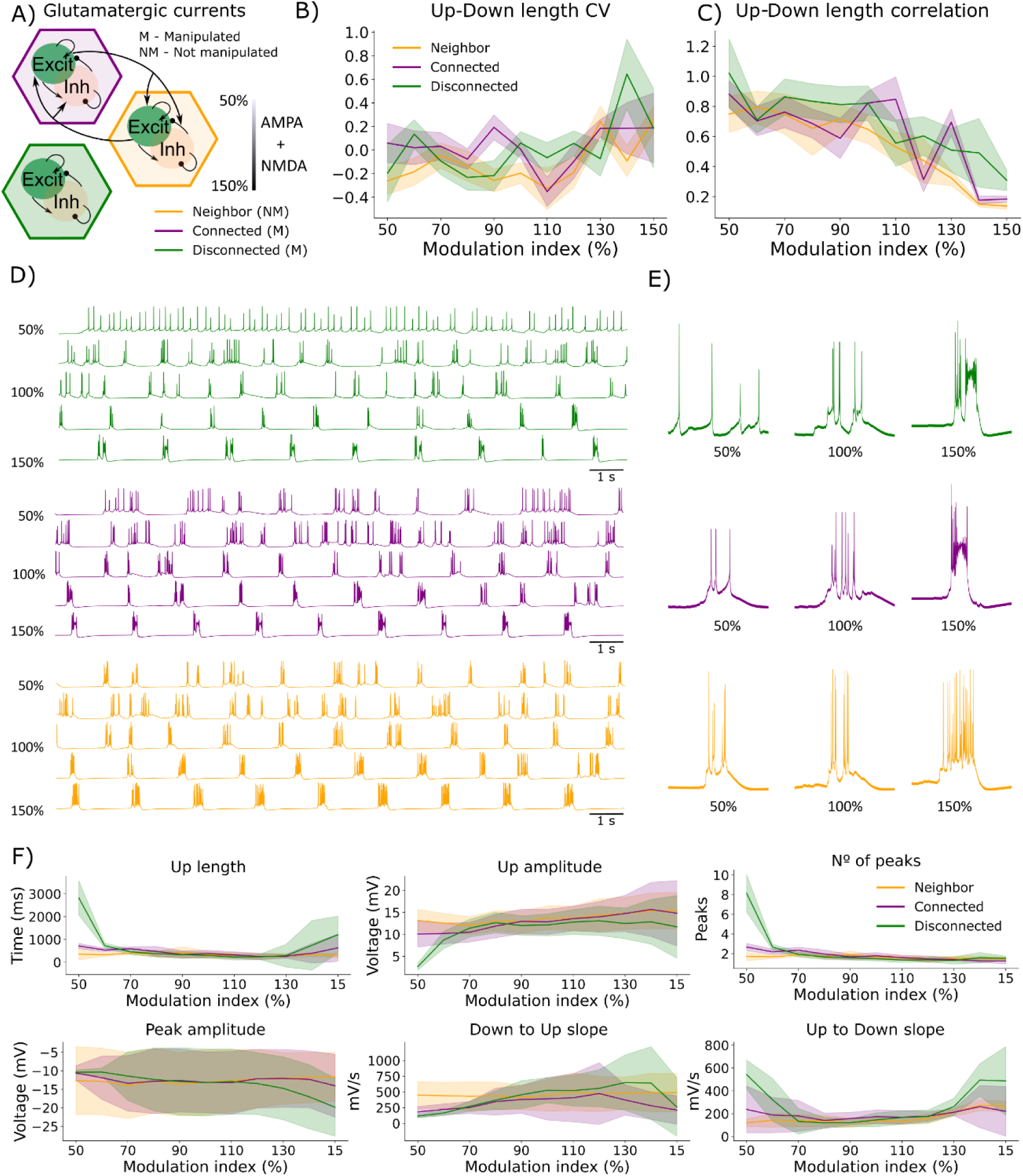
Impact of the manipulation of glutamatergic conductance on SWA. **A)** Schematic representation of the simulations **B)** Coefficient of variance of the cycle length for the different levels of manipulation. **C)** Correlation of the length of each Up state and the following Down state for the different levels of manipulation. **D)** Representative traces of the neurons in the three manipulations shown in A) during different levels of manipulation (from 50% to 150%). **E)** Up states examples of the traces in D). Each trace is 1 second long. **F)** Impact on the properties of the Up states resulted from the simulations shown in A).

The quantification of the Up state parameters (Fig. 6f) showed marked differences between the simulated populations. Overall, the manipulation of glutamatergic drive was correlated with the length and amplitude of the Up states, the number of peaks and the speed of the slopes between transitions. On the other hand, peak amplitude was not altered in the connected populations, and inversely related to the modulation of the glutamatergic drive in the disconnected population. Manipulations that decreased glutamatergic drive led to local changes. In particular, the length and number of peaks of the Up states, as well as the Down to Up slope decreased more acutely in the manipulated population than in its neighbor. This is also true in a smaller scale for the Up to Down transition. At last, the strongest increment in glutamatergic conductance we explored (150%) pushed the manipulated populations into a regime that resembled ictal activity recorded by intracellular electrodes (Koerling et al. 2019; Winograd, Destexhe, and Sanchez-Vives 2008), in which strong depolarizations inactivate action potentials (Fig. 5c); from now on we will refer to it as epileptiform-like activity. This came together with an increment of firing rate in the neighbor population (Fig. 5b,c), albeit without being pushed to this network state.

In general, modulation in glutamatergic drive has strong impact in the SWA, reduction or increments of 50% of the modulation index transform the SWA in an asynchronous activity (or close to) up to highly synchronous state, resembling epileptiform activity, respectively.

### Balanced contribution of AMPA and NMDA to the properties of the Up states

To obtain a finer grained picture, we studied the different contribution of the glutamate driven synaptic currents in the model to the Up state properties, as well as their impact on local and global dynamics. To understand this, we run a new set of simulations of two networks as in Fig. 6, but now only AMPA (Fig. 7a-d) or NMDA (Fig. 7 e-h) currents were modified in a range of 50% to 150% of the original conductance (Fig. 7). This gave a finer grained understanding of the relative contribution of each synaptic current to the SWA.

**Fig. 7.**
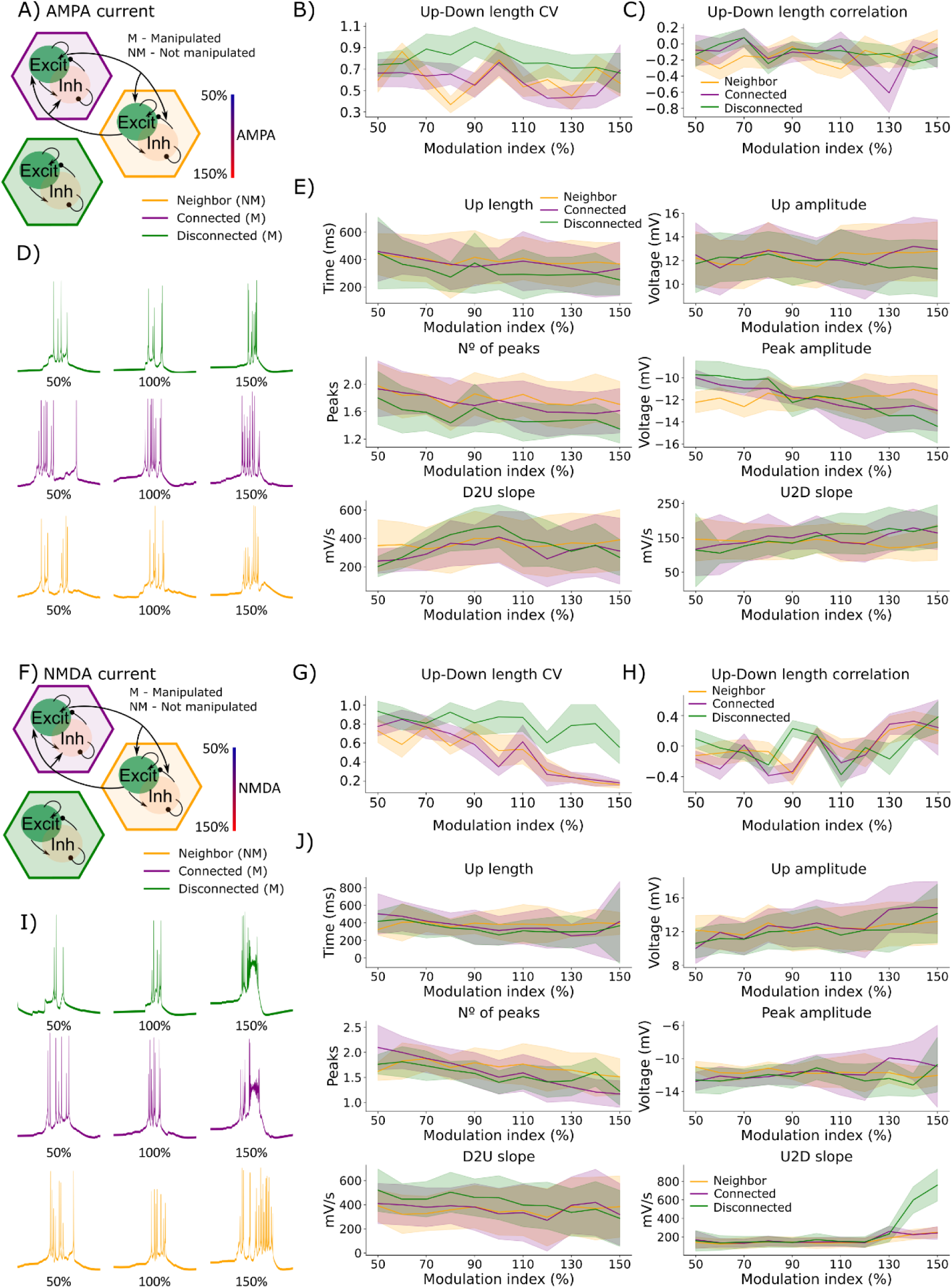
Impact of the manipulation of AMPA and NMDA conductance on SWA. **A)** Schematic representation of the simulations of AMPA conductance. **B)** Coefficient of variance of the cycle length for the different levels of manipulation. **C)** Correlation of the length of each Up state and the following Down state for the different levels of manipulation (50-150%). **D)** Examples of Up states of neurons from the simulated populations shown in A), for different levels of manipulation. Each trace is 1 second long. **E)** Impact on the properties of the Up states of the manipulation shown in A). **F-J)** Equivalent to A-E, simulating the manipulation of NMDA conductance.

Interestingly, the impact of glutamatergic manipulations in the cycle CV and Up and Down length correlation (Fig. 6b,c) was not reproduced when AMPA was manipulated independently. Changes in AMPA had no impact in those parameters (Fig. 7b,c), while NMDA caused results equivalent to the manipulations performed in Fig. 6, decreasing the CV and increasing the Up and Down length correlation within each cycle.

Then we explored the impact of the manipulation of AMPA and NMDA currents in the Up states (examples in Fig. 7e,j and e). Most of them has similar effects, however epileptiform-like activity was reproduced only in the manipulated populations in which NMDA was increased up to 150%, showing that NMDA currents are sufficient to explain this dynamical state, previously shown in Fig. 6d,e. The level of manipulation was in both cases inversely correlated with the length and number of peaks of the Up states, without a relevant impact in their amplitude and the speed of the Up to Down transition, except for an acceleration for the strongest values of the manipulation of the NMDA current, reproducing the result in Fig. 6f. Peak amplitude slightly increased with the strength of both currents. The increase observed in the Down to Up slope, when glutamatergic modulation was augmented (Fig. 6) was not reproduced manipulating either AMPA or NMDA, suggesting that changes in both currents are necessary. We found that the impact on this parameter of one of the currents had an opposite effect to the manipulation of AMPA and NMDA together, leading to slower slopes as either current conductance increased.

### GABAergic currents

At last, we explored the impact of manipulating GABAergic drive (Fig. 8a). This led to changes in the model in the opposite direction to glutamate, but with smaller magnitude, especially in the case of the biggest perturbations (50% or 150% drive). The CV of the cycle’s length was directly related to the strength of the manipulation, with more variable cycles as GABAergic drive increased (Fig. 8b), and a reduced correlation between the length of each state (Fig.8c). Examples showing the impact of the manipulation in the output of the model are shown in Fig. 8 d and e. On the contrary to Fig. 6, GABAergic manipulations decreasing the conductance up to a 50% never pushed the system into an asynchronous state, suggesting that, at least during the strongly depolarized Up states, glutamatergic synapsis may exert a stiffer control on the global dynamics than inhibitory ones, probably because of the compensatory contribution of activity dependent K+ channels.

**Fig. 8.**
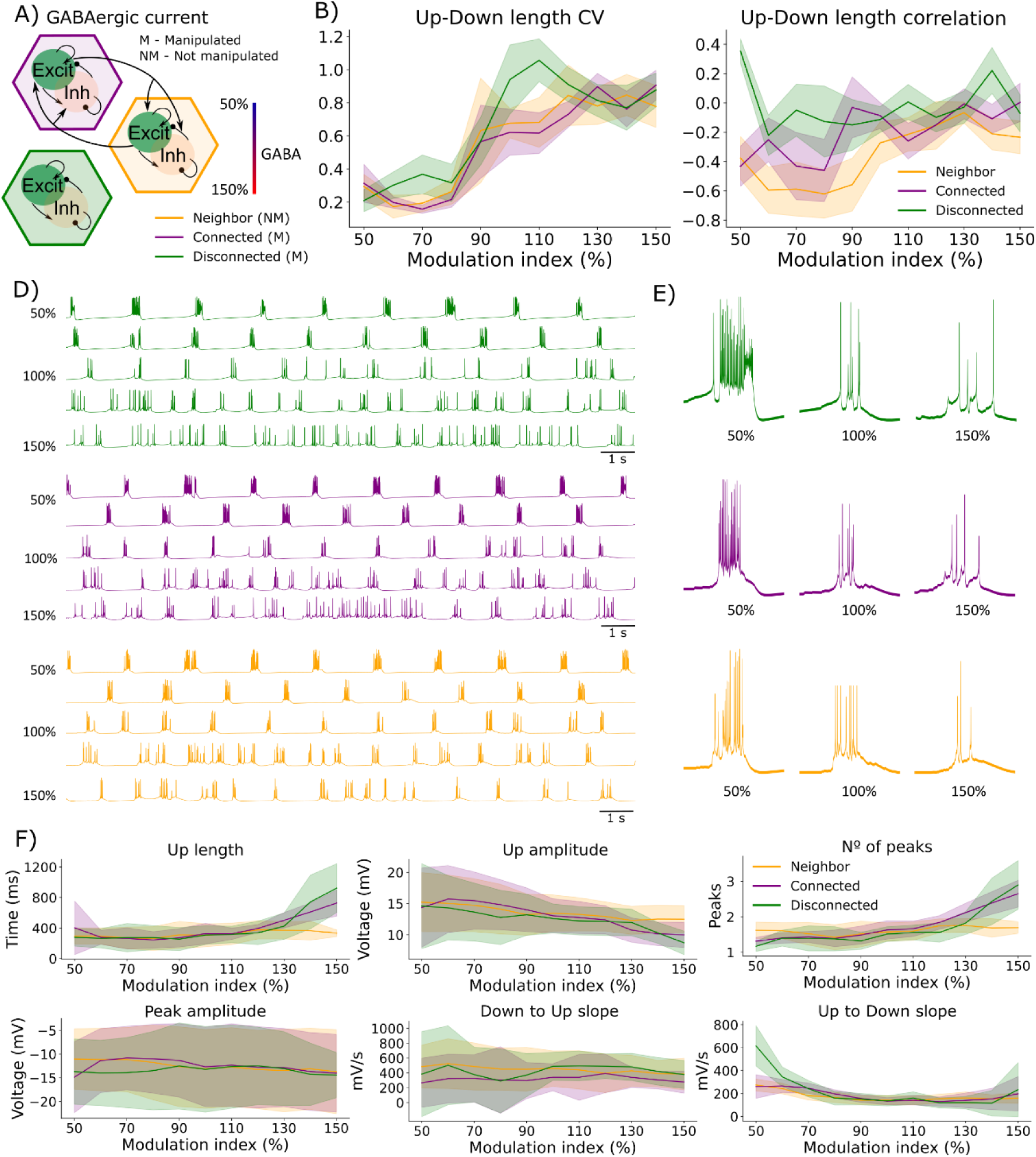
Impact of the manipulation of GABAergic conductance on SWA. **A)** Schematic representation of the simulations **B)** Coefficient of variance of the cycle length for the different levels of manipulation. **C)** Correlation of the length of each Up state and the following Down state for the different levels of manipulation. **D)** Representative traces of the neurons in the three manipulations shown in A) during different levels of manipulation (from 50% to 150%). **E)** Up states examples of the traces in D). Each trace is 1 second long. **F)** Impact on the properties of the Up states resulted from the simulations shown in A).

Despite a clear rise in firing rate (Fig. 8d,e weakening GABAergic drive, up to 50%, had a moderate impact in the Up state properties (Fig. 8f). It was also related with an increase in the amplitude and the Up to Down slope, as previously described on ex vivo slices (Sanchez-Vives et al. 2010), the later particularly strong in the disconnected population. A modest reduction in the length and number of peaks of the Up state was also observed. Increasing the GABAergic drive was related to longer Up states with more peaks and smaller amplitude of the manipulated populations. Note how very small changes are detected in the Up state properties of the neighbor population (Fig. 8f).

## Discussion

The SWA regime has been considered the default state of the cortical network (Sanchez-Vives, Massimini, and Mattia 2017). It is a robust low-complexity activity pattern on which cortical populations transition from quiescent Down to active Up states, which propagate to their postsynaptic targets, creating travelling waves (Massimini et al. 2004; Murphy et al. 2009; Pazienti et al. 2022). While cortical populations transition among states at a similar frequency and the properties of the Up states are remarkably similar across cortical regions, some differences have already been described (Ruiz-Mejias et al. 2011; Alegre-Cortés et al. 2021). This makes the SWA and ideal regime of brain activity to explore how the local features and long-range inputs of a circuit determine its dynamics. To do so, we generated a conductance-based model of cortical Layer V populations under the SWA and organized our work around two main scientific questions. First, we have explored the influence of long-range connections between populations. For this purpose, we built populations with different levels of excitability by means of manipulating their leak current (Fig.1) and induced a homogeneous frequency of Up states introducing long-range projections in the model (Fig. 2). We have also shown *in vivo* how the preferred axis of propagation of slow waves can be explained by means of asymmetries in the excitability of different populations (Fig.3). Then, we explored how long-range projections impacted the properties of the Up states and tended to homogenize them, blurring local differences (Fig. 4). Second, we performed a sensitivity analysis, exploring how manipulations of different cellular and synaptic components of the circuit impacted in the Up states (Fig. 5-8).

It is relevant to explore how does the system output change following the manipulation of its key components and how certain activity patterns can emerge from different combination of parameters (Marder and Goaillard 2006; Prinz, Bucher, and Marder 2004). We argue that Up states are an open window to explore stiff manifolds in the parameter space of cortical circuits (Gutenkunst et al. 2007). Here we show that these stiff manifolds are associated to coordinated changes in key cellular and synaptic properties that, being resilient to the influence of long-range inputs, can effectively shape SWA and can be inferred from experimentally recorded activity.

A relevant example of a local feature was the number of peaks in the up state that determines the presence of diplets (or potentially triplets, quadruplets…). They are sharp V-shaped deflections of the membrane voltage that occur in vivo spontaneously during the Up state, giving them a particular horned-like shape, and are visible in some cortical and subcortical regions, including at the level of the LFP. Because during the valley of the diplets, the membrane voltage moves far from the spiking threshold, they constrain the possible firing patterns of the neurons during the Up state. They do occur locally in different cortical areas involved in movement (M1, S1, FrA) as well as at least one subcortical region, the dorsolateral striatum (Alegre-Cortés et al. 2021). From a system-dynamics perspective, the occurrence of quasi-periodic bursts of activity is very likely associated with a collective phenomenon where the population firing rate displays damped oscillations. This happens if the observed cortical area is close to a so-called “Hopf bifurcation” (Brunel and Hakim 1999; Bressloff and Coombes 2000; Linaro, Storace, and Mattia 2011). In this case the cortical network is trapped into a stable focus approaching an equilibrium point via fading out of oscillations. These damped oscillations can be elicited by the intrinsic fluctuations of their firing rate since cortical circuits are composed of a finite number of neurons (Vinci, Benzi, and Mattia 2023). Persistence of oscillations is determined by the distance from the critical point, such that closer is the network to this point (i.e., in the parameter space shown Fig. 9A) more oscillations will be visible following the Up-state onset. In this framework cortical areas showing diplets or triplets are those closest to display the mentioned critical dynamics. The predominant presence of these damped oscillations in movement related areas makes then diplets an interesting target to study, as they may be the reflection of some shared circuit constraint that shape the neural code of these brain regions.

**Fig. 9.**
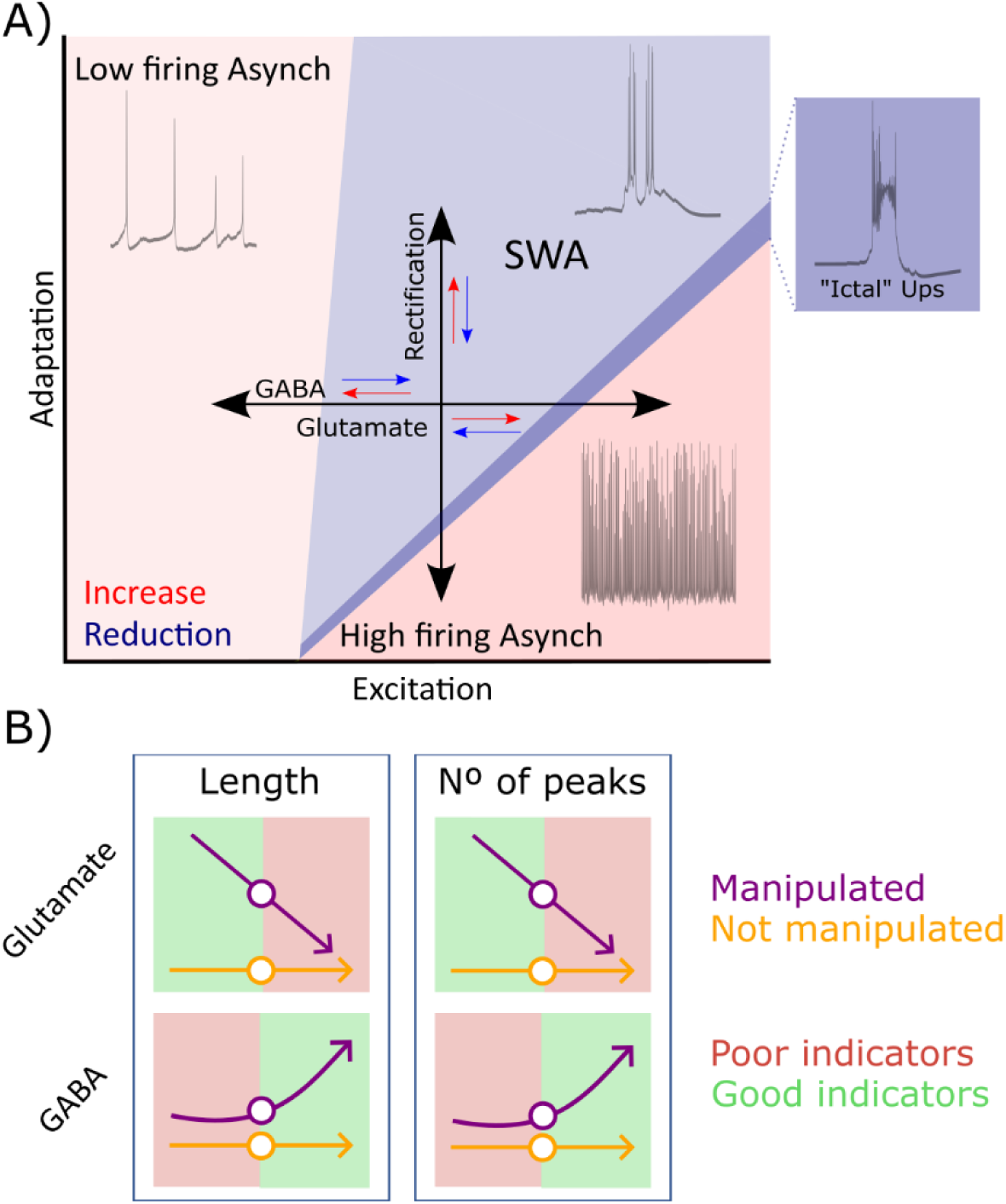
SWA mechanisms behind global and local motifs. **A)** Sketch representing the main mechanisms that modulate the SWA and the transition to other dynamical regimes, depicted in different colors. Darker blue represents a parameter space on which ictal Up state alternate with Down states, in a pre-epileptic state (bifurcation diagram adapted from Mattia and Sanchez-Vives 2012). **B)** Summary of the range of manipulations that lead to local changes in the Up states, from 50 to 150%. Green represents the manipulation range on which changes can be detected only in the connected population (purple) and not in its connected neighbor (orange).

### SWA sources in vivo and in silico

When a cortical population transitions into an Up state, this one can be self-generated or propagated from another population. (Pazienti et al. 2022). This can explain why when a cortical slab is deafferented it can still create Up states, but the frequency of occurrence is lower than in the neighboring tissue (Timofeev 2000) and how even very small ex vivo cortical slices (<2 x 5 mm) can generate slow waves (Sanchez-Vives et al. 2007). We theorized that in a scenario of connected populations, the one of higher excitability would impose its own SWA rhythm to the others, acting as the leader of the waves. In this work, we challenged this assumption both *in silico* and *in vivo*. We first showed how the frequency of the Up states of an isolated population can be modulated by its excitability level, manipulated by means of the conductance of the Cl-leak current in our computational model (Compte et al. 2003). On the other hand, when populations our coupled following biologically plausible connection probabilities (Potjans and Diesmann 2014), the least excitable population becomes a follower of the other, having the same number of Up states. A specific prediction of the model is that the cortical frequency of oscillation is modulated by a leading population, and not because of multiple populations of lower intrinsic frequency connected. Thus, we predict the presence of “hot spots”, populations that can potentially oscillate at a higher frequency than others in the cortex and lead and impose their own rhythm to the cortex. Consistent with our results, (Ruiz-Mejias et al. 2011) shown a homogeneous number of Up states along the cortex, with a tendency to initiate in the prefrontal cortex, which displayed a slightly higher firing rate, in line with the results shown in Fig. 2.

Together with the homogenization of the SWA frequency, coupling populations led to a regime of a lower dimensionality, in line with previous results using recurrent neuronal networks (Clark and Beiran 2024; Kitano and Fukai 2007). This is in line with what expected in network of coupled oscillators (Strogatz 2000; Kuramoto 1984). Furthermore, the firing rate of both populations increased and became closer to the one described *in vivo* (Yousheng, Hasenstaub, and McCormick 2003; El Boustani et al. 2007) without further modifications of the cellular or circuit properties of the model, as well as an increment in the length of the Up state and the speed of the transition to the Up state, and the number of peaks, as previously shown to be caused by a stronger recruitment of outward rectification (Compte et al. 2003).

We then tested our prediction *in vivo*, increasing the local excitability of the caudal cortical pole releasing NE and CCh (Schmidt et al. 2013; D’Andola et al. 2018; Foehring, Schwindt, and Crill 1989). This led to an increase in the frequency of SWA around the area of release, which we computed based on the logMUA. This in turn shifted the preferential directionality of the SWA from rostrocaudal to caudorostral and imposed a new frequency of Up states in the cortex. This opens the question about the cause behind the original rostrocaudal directionality. This may be caused by several mechanisms, including a higher level of intrinsic excitability of the prefrontal cortex as the one included in the model, potentially mediated by its dense cholinergic innervation (Bloem, Poorthuis, and Mansvelder 2014) that can enhance neuronal excitability (Picciotto, Higley, and Mineur 2012), or by the dopamine present in the prefrontal cortex during sleep (Monti and Monti 2007), acting through the D1 receptors present in pyramidal and VIP+ interneurons (Anastasiades, Boada, and Carter 2019). Alternatively, a higher level of excitability can be potentially driven by the increased level of excitatory recurrence of connections in prefrontal cortex (Y. Wang et al. 2006; Cakan et al. 2022) or by the contribution of presynaptic inputs; i.e., claustrum has been widely considered to be involved in the sleep-wake cycle (Narikiyo et al. 2020) and Slow Waves emerge at the age on which claustral projections of the anterior cingulate cortex maturate (Hoerder-Suabedissen et al. 2023). Another possible relevant input comes from the olfactory bulb, given the role of breathing in the modulation of the Up to Down transition (Karalis and Sirota 2022).

### Sloppy components underlie robust cortical computations

Up states are the result of a high level of balanced excitatory and inhibitory synaptic activity together with cellular adaptation mechanisms. And excess of inhibition can abolish the emergence of spontaneous Up states, while a disbalance towards excessive excitation can push the system into an epileptic state (Koerling et al. 2019; Sanchez-Vives et al. 2010). Most of our understanding of the key components that underlie SWA come from *ex vivo* perturbations, and the computational models they have guided. Despite this extensive body of knowledge, it is still unclear how to explain the differences between the Up states recorded in connected cortical areas, as similar features can result from changes in different components of the circuit (Gutenkunst et al. 2007; Prinz, Bucher, and Marder 2004; Marder and Goaillard 2006; Schulz, Goaillard, and Marder 2006). In this context, we have performed a sensitivity analysis of different circuit components that are relevant for the SWA (Figs. 5-8), exploring how they are affected by the presence of long-range inputs and detecting which changes had a global effect on the network or on the contrary only altered the dynamics of the manipulated population. First, we showed that the manipulation of rectifying currents (Fig. 5) had an asymmetric effect, increasing the conductance of these currents had a minor effect in the properties of the Up states (Fig. 5f), while when they were reduced, a global increment of the Up state length and number of peaks, together with a decrease in their amplitude. The manipulation of glutamatergic (Fig. 6,7) and GABAergic (Fig. 8) synaptic currents had opposed effects (Fig. 9a). Decreasing glutamate or increasing GABA led to longer Up states with a higher number of peaks. These changes were mostly local in connected populations, suggesting that these features are relevant to exploring the local excitation/inhibition ratio of a population (Fig. 9b). On the other hand, increases in glutamate and decreases in GABA had a smaller effect on the subthreshold properties of the Up state, and in the wider range of manipulation (Figs 6,8 d,e) pushed the populations to a (pre)epileptic state (Fig. 9a) similar to equivalent perturbations performed in *ex vivo* (Koerling et al. 2019; Sanchez-Vives et al. 2010), with a similar impact on the nonmanipulated neighbor. Note how the suppression of spikes caused by the strong increase of glutamatergic currents, NMDA in particular (Fig. 7), reproduced the ictal Up states in (Koerling et al. 2019; Sanchez-Vives et al. 2008). Interestingly, we did not observe local changes in the cycle CV (Figs 5-8); whenever a manipulation affected this property, the changes were extended to the neighbor population, consistent with the homogeneity in cycle properties seen in vivo (Ruiz-Mejias et al. 2011). We will briefly discuss the physiological relevance of the main results from the sensitivity analysis in the following sections.

### Disparate modulation of SWA by synaptic or intrinsic hyperpolarizing currents

Low dimensional models of SWA clearly distinguish between the effect of inhibition, short-term plasticity and adaptation (Parga and Abbott 2007; Sanchez-Vives, Massimini, and Mattia 2017; Torao-Angosto et al. 2021). Despite this theoretical understanding, it is challenging to detect in vivo whether differences in the recruitment of hyperpolarizing currents are caused due to synaptic (GABA mediated) or intrinsic channels (rectification mediated). We have shown how small (up to +/-50%) of these currents have opposing effects in the Up state and cycle properties of the SWA (Figs 5, 8). We have also shown how changes in the GABAergic currents had a local effect, longer Up states with more peaks when GABA conductance was increased, that are not seen in the simulations on which outward rectifiers were manipulated.

We expect that our results will guide futures experiments aiming to decipher the mechanism underlying the hyperpolarizing sources of distinct cortical circuits, through the comparison of their SWA.

### Mechanistic insights obtained from the analysis of diplets

Diplets are one of the most relevant differences between the Up states of somatosensory and motor cortical areas compared to other cortical regions (Alegre-Cortés et al. 2021), and have a major impact on the subthreshold and firing patterns of neurons during SWA regime. They are also preserved across other subcortical motor related areas, such as the dorsolateral striatum. Nevertheless, the mechanistic origin of diplets, whether *in vivo* or not, remains an open question. We found different possible mechanisms that caused an increase in the number of diplets in our simulations, both locally and globally. Local diplets were obtained in the manipulations that lead to a decrease of Glutamatergic force (Fig. 6) or an increase of GABAergic conductance (Fig. 8), suggesting that this can be a useful activity motif to explore the synaptic E/I balance of a circuit. In the case of the manipulation of glutamatergic drive, it was necessary to manipulate AMPA and NMDA currents together to generate a local increase in the number of peaks (Fig. 7). An alternative mechanism that led to an increase in the number of peaks in the Up state was mediated by the reduction of Na^+^ and Ca^2+^ activated K^+^ channels, but in the case the same effect was found in the neighbor population. Summarizing, we have described how changes in the main agents that modulate SWA can be the substrate of diplets inside the Up state.

Lastly, it is relevant to comment a possible limitation of our work, as our simulations decreasing GABAergic conductance did not reproduce previous results, showing the emergence of diplets following bicuculline administration (Sanchez-Vives et al. 2010). There are two possible explanations for this: First, we have restricted our sensitivity analysis to a range of manipulations (50%-150%) that could be considered within physiological range, and not the result of strong pharmacological perturbations. Thus, it may be the case that the reduction of GABAergic conductance mediated by bicuculline in this work was stronger than the range we explored. Second, bicuculline blocks not just GABAergic receptors but also Ca^2+^ activated K^+^ channels (Khawaled et al. 1999). Given that we show an increase in the number of peaks following the reductions of rectification currents (Fig. 5), this is a possible explanation of the diplets shown in (Sanchez-Vives et al. 2010).

### Transition from SWA to epileptic activity

Some types of cortical epilepsies occur with a higher frequency during NREM sleep, on which SWA can be recorded (Koerling et al. 2019; Bazil and Walczak 1997; Crespel, Baldy-Moulinier, and Coubes 1998), suggesting a possible pro-epileptogenic property of this brain regime when the fine balance of excitation and inhibition that governs the Up states is altered (Fig. 9). In this context, our sensitivity analysis shows how Up states are transformed into ictal activity by a significant increase of glutamatergic force (Figs. 6,7) or decrease of inhibition (Fig. 8). Manipulations of an equivalent magnitude of outward rectifying currents did not lead to pre-epileptic Up states.

We expect that our work can be useful in the study of (pre)epileptic networks, either pointing at Up state properties of regions that are susceptible to become epileptic, or through the impact of epileptic Up states into neighbor populations. A missing link remains that can explain the ease of the transition from SWA to some types of epileptic states. A mechanism of potential interest is through ATP mediated K+ currents (Inagaki et al. 1995; 1996), as they can modulate both the SWA activity (Cunningham et al. 2006) and, at least, certain types of epilepsies (Eydam, Franović, and Kang 2024; Kovács et al. 2018; Bazzigaluppi et al. 2017)

## Conclusions

In summary, using our model we have explored how global and local components of a circuit shape its SWA, frequency of Up states and axis of propagation. Following our sensitivity analysis, we have reported several Up state properties than can be useful to seek for differences between cortical circuits, as well as other features that are sensitive to the status of the whole network and therefore poor features to study circuits locally. We belief our results will be a useful substrate to guide future research beyond the use of SWA to interrogate circuit properties. This includes the attraction into the SWA of neighbor regions that would not normally be in this dynamic regime (Massimini et al. 2024; Sarasso et al. 2020) and the characterization of the transition from SWA to epileptic states (Bazil and Walczak 1997; Crespel, Baldy-Moulinier, and Coubes 1998).

## Acknowledgements

This work is supported by the Severo Ochoa Grant CEX2021-001165-S, funded by MCIN/AEI/ 10.13039/501100011033; PID2021-129070NB-I00 funded by MCIN and Parkinson NEURO-AGING+; Parkinson. NEURO-AGING+ Ref. RECEUNAG2021; MULTISYN, Ref. CIPROM/2022/08, Generalitat Valenciana. JAC is supported by Ministerio de Universidad, European Union and Universidad Miguel Hernández de Elche through a Margarita Salas fellowship. MM is supported by funding from the Italian National Recovery and Resilience Plan (PNRR), M4C2, funded by the European Union–NextGenerationEU (Project IR0000011, CUP B51E22000150006, “EBRAINS-Italy”). MS is supported by Ministerio de Ciencia e Innovación “Ayudas Juan de la Cierva-formación” FJC2021-047618-I grant.

## Bibliography

Alegre-Cortés, J., M. Sáez, R. Montanari, and Ramón Reig. 2021. ‘Medium Spiny Neurons Activity Reveals the Discrete Segregation of Mouse Dorsal Striatum’. ELife 10 (February):1–41. 10.7554/eLife.60580.

Anastasiades, Paul G., Christina Boada, and Adam G. Carter. 2019. ‘Cell-Type-Specific D1 Dopamine Receptor Modulation of Projection Neurons and Interneurons in the Prefrontal Cortex’. Cerebral Cortex 29 (7): 3224–42. 10.1093/cercor/bhy299.

Barbero-Castillo, Almudena, Pedro Mateos-Aparicio, Leonardo Dalla Porta, Alessandra Camassa, Lorena Perez-Mendez, and Maria V. Sanchez-Vives. 2021. ‘Impact of Gabaa and Gabab Inhibition on Cortical Dynamics and Perturbational Complexity during Synchronous and Desynchronized States’. Journal of Neuroscience 41 (23): 5029–44. 10.1523/JNEUROSCI.1837-20.2021.

Bazil, Carl W., and Thaddeus S. Walczak. 1997. ‘Effects of Sleep and Sleep Stage on Epileptic and Nonepileptic Seizures’. Epilepsia 38 (1): 56–62. 10.1111/j.1528-1157.1997.tb01077.x.

Bazzigaluppi, Paolo, Azin Ebrahim Amini, Iliya Weisspapir, Bojana Stefanovic, and Peter L. Carlen. 2017. ‘Hungry Neurons: Metabolic Insights on Seizure Dynamics’. International Journal of Molecular Sciences. MDPI AG. 10.3390/ijms18112269.

Beltramo, Riccardo, Giulia D’Urso, Marco Dal Maschio, Pasqualina Farisello, Serena Bovetti, Yoanne Clovis, Glenda Lassi, Valter Tucci, Davide De Pietri Tonelli, and Tommaso Fellin. 2013. ‘Layer-Specific Excitatory Circuits Differentially Control Recurrent Network Dynamics in the Neocortex’. Nature Neuroscience 16 (2): 227–34. 10.1038/nn.3306.

Berger, H. 1929. ‘Uber Das Elektrenkephalogramm Des Menschen.’ J. Psycho. Neurol. 40 (1875): 160–79.

Bloem, Bernard, Rogier B. Poorthuis, and Huibert D. Mansvelder. 2014. ‘Cholinergic Modulation of the Medial Prefrontal Cortex: The Role of Nicotinic Receptors in Attention and Regulation of Neuronal Activity’. Frontiers in Neural Circuits. Frontiers Research Foundation. 10.3389/fncir.2014.00017.

Boustani, Sami El, Martin Pospischil, Michelle Rudolph-Lilith, and Alain Destexhe. 2007. ‘Activated Cortical States: Experiments, Analyses and Models’. Journal of Physiology 101 (1–3): 99–109. 10.1016/j.jphysparis.2007.10.001.

Bressloff, Paul C, and S Coombes. 2000. ‘Dynamics of Strongly Coupled Spiking Neurons’. Neural Computation, no. 12, 91–129.

Brunel, Nicolas, and Vincent Hakim. 1999. ‘Fast Global Oscillations in Networks of Integrate-and-Fire Neurons with Low Firing Rates’. Neural Computation, no. 11, 1621–71.

Cakan, Caglar, Cristiana Dimulescu, Liliia Khakimova, Daniela Obst, Agnes Flöel, and Klaus Obermayer. 2022. ‘Spatiotemporal Patterns of Adaptation-Induced Slow Oscillations in a Whole-Brain Model of Slow-Wave Sleep’. Frontiers in Computational Neuroscience 15 (January): 1–19. 10.3389/fncom.2021.800101.

Capone, Cristiano, Beatriz Rebollo, Alberto Munoz, Xavi Illa, Paolo Del Giudice, Maria V. Sanchez-Vives, and Maurizio Mattia. 2019. ‘Slow Waves in Cortical Slices: How Spontaneous Activity Is Shaped by Laminar Structure’. Cerebral Cortex 29 (1): 319–35. 10.1093/cercor/bhx326.

Clark, David G., and Manuel Beiran. 2024. ‘Structure of Activity in Multiregion Recurrent Neural Networks’. ArXiv, February. http://arxiv.org/abs/2402.12188.

Compte, Albert, Maria V. Sanchez-Vives, David A. McCormick, and Xiao Jing Wang. 2003. ‘Cellular and Network Mechanisms of Slow Oscillatory Activity (<1 Hz) and Wave Propagations in a Cortical Network Model’. Journal of Neurophysiology 89 (5): 2707–25. 10.1152/jn.00845.2002.

Cowan, R. L., and C. J. Wilson. 1994. ‘Spontaneous Firing Patterns and Axonal Projections of Single Corticostriatal Neurons in the Rat Medial Agranular Cortex’. Journal of Neurophysiology 71 (1): 17–32. 10.1152/jn.1994.71.1.17.

Crespel, Arielle, Michel Baldy-Moulinier, and Philippe Coubes. 1998. ‘The Relationship between Sleep and Epilepsy in Frontal and Temporal Lobe Epilepsies: Practical and Physiopathologic Considerations’. Epilepsia 39 (2): 150–57. 10.1111/j.1528-1157.1998.tb01352.x.

Cunningham, M O, D D Pervouchine, C Racca, N J Kopell, C H Davies, R S G Jones, R D Traub, and M A Whittington. 2006. ‘Neuronal Metabolism Governs Cortical Network Response State’. Proceedings of the National Academy of Sciences of the United States of America 103 (14): 5597–5601. doi www.pnas.orgcgi10.1073pnas.0600604103.

Dalla Porta, Leonardo, Almudena Barbero-Castillo, José Manuel Sanchez-Sanchez, Nathalia Cancino, and Maria V Sanchez-Vives. 2024. ‘H-Current Modulation of Cortical Up and Down States’. BioRxiv. 10.1101/2024.04.05.588281.

Dalla Porta, Leonardo, Almudena Barbero-Castillo, Jose Manuel Sanchez-Sanchez, and Maria V. Sanchez-Vives. 2023. ‘M-Current Modulation of Cortical Slow Oscillations: Network Dynamics and Computational Modeling’. PLoS Computational Biology 19 (7): 1–17. 10.1371/journal.pcbi.1011246.

D’Andola, Mattia, Beatriz Rebollo, Adenauer G. Casali, Julia F. Weinert, Andrea Pigorini, Rosa Villa, Marcello Massimini, and Maria V. Sanchez-Vives. 2018. ‘Bistability, Causality, and Complexity in Cortical Networks: An in Vitro Perturbational Study’. Cerebral Cortex 28 (7): 2233–42. 10.1093/cercor/bhx122.

Dasilva, Miguel, Alessandra Camassa, Alvaro Navarro-Guzman, Antonio Pazienti, Lorena Perez-Mendez, Gorka Zamora-López, Maurizio Mattia, and Maria V. Sanchez-Vives. 2021. ‘Modulation of Cortical Slow Oscillations and Complexity across Anesthesia Levels’. NeuroImage 224:117415. 10.1016/j.neuroimage.2020.117415.

Destexhe, Alain, Diego Contreras, and Mircea Steriade. 1999. ‘Spatiotemporal Analysis of Local Field Potentials and Unit Discharges in Cat Cerebral Cortex during Natural Wake and Sleep States’. Journal of Neuroscience 19 (11): 4595– 4608. 10.1523/jneurosci.19-11-04595.1999.

Eydam, Sebastian, Igor Franović, and Louis Kang. 2024. ‘Control of Seizure-like Dynamics in Neuronal Populations with Excitability Adaptation Related to Ketogenic Diet’. Chaos 34 (5). 10.1063/5.0180954.

Foehring, R C, P C Schwindt, and W E Crill. 1989. ‘Norepinephrine Selectively Reduces Slow Ca2+-and Na+-Mediated K+ Currents in Cat Neocortical Neurons’. Journal of Neurophysiology 61 (2): 245–56. www.physiology.org/journal/jn.

Gigante, Guido, Maurizio Mattia, and Paolo Del Giudice. 2007. ‘Diverse Population-Bursting Modes of Adapting Spiking Neurons’. Physical Review Letters 98 (14). 10.1103/PhysRevLett.98.148101.

Greenberg, Anastasia, Javad Karimi Abadchi, Clayton T. Dickson, and Majid H. Mohajerani. 2018. ‘New Waves: Rhythmic Electrical Field Stimulation Systematically Alters Spontaneous Slow Dynamics across Mouse Neocortex’. NeuroImage 174 (July):328–39. 10.1016/j.neuroimage.2018.03.019.

Gutenkunst, Ryan N., Joshua J. Waterfall, Fergal P. Casey, Kevin S. Brown, Christopher R. Myers, and James P. Sethna. 2007. ‘Universally Sloppy Parameter Sensitivities in Systems Biology Models’. PLoS Computational Biology 3 (10): 1871–78. 10.1371/journal.pcbi.0030189.

Haider, Bilal, and David A. McCormick. 2009. ‘Rapid Neocortical Dynamics: Cellular and Network Mechanisms’. Neuron 62 (2): 171–89. 10.1016/j.neuron.2009.04.008.Rapid.

Hoerder-Suabedissen, Anna, Gabriel Ocana-Santero, Thomas H. Draper, Sophie A. Scott, Jesse G. Kimani, Andrew M. Shelton, Simon J.B. Butt, Zoltán Molnár, and Adam M. Packer. 2023. ‘Temporal Origin of Mouse Claustrum and Development of Its Cortical Projections’. Cerebral Cortex 33 (7): 3944–59. 10.1093/cercor/bhac318.

Idesis, Sebastian, Gustavo Patow, Michele Allegra, Jakub Vohryzek, Yonatan Sanz Perl, Maria V. Sanchez-Vives, Marcello Massimini, Maurizio Corbetta, and Gustavo Deco. 2024. ‘Whole-Brain Model Replicates Sleep-like Slow-Wave Dynamics Generated by Stroke Lesions’. Neurobiology of Disease 200 (October). 10.1016/j.nbd.2024.106613.

Inagaki, Nobuya, Tohru Gonoi, John P Clement IV, Noriyuki Namba, Johji Inazawa, Gabriela Gonzalez, Aguilar-Bryan Lydia, Susumu Seino, and Joseph Bryan. 1995. ‘Reconsttitution of IKATP: An Inward Rectifier Subunit Plos the Sulfonyurea Receptor’. Science 270 (November):1166–70. DOI:10.1126/science.270.5239.1166.

Inagaki, Nobuya, Tohru Gonoi, John P Clement, Chang-Zheng Wang, Lydia Aguilar-Bryan, Joseph Bryan, and Susumu Seino. 1996. ‘A Family of Sulfonylurea Receptors Determines the Pharmacological Properties of ATP-Sensitive K Channels’. Neuron 16 (May):1011–17.

Jercog, Daniel, Alex Roxin, Peter Barthó, Artur Luczak, Albert Compte, and Jaime de la Rocha. 2017. ‘UP-DOWN Cortical Dynamics Reflect State Transitions in a Bistable Network’. *ELife*, August. 10.7554/eLife.22425.001.

Karalis, Nikolaos, and Anton Sirota. 2022. ‘Breathing Coordinates Cortico-Hippocampal Dynamics in Mice during Offline States’. Nature Communications 13 (1). 10.1038/s41467-022-28090-5.

Khawaled, Radwan, Andrew Bruening-Wright, John P. Adelman, and James Maylie. 1999. ‘Bicuculline Block of Small-Conductance Calcium-Activated Potassium Channels’. European Journal of Physiology 438 (3): 314–21. 10.1007/s004240050915.

Kitano, Katsunori, and Tomoki Fukai. 2007. ‘Variability v.s. Synchronicity of Neuronal Activity in Local Cortical Network Models with Different Wiring Topologies’. Journal of Computational Neuroscience 23 (2): 237–50. 10.1007/s10827-007-0030-1.

Koerling, Anna Lucia, Tanja Fuchsberger, Ole Paulsen, and Y. Audrey Hay. 2019. ‘Partial Restoration of Physiological UP-State Activity by GABA Pathway Modulation in an Acute Brain Slice Model of Epilepsy’. Neuropharmacology 148 (April):394–405. 10.1016/j.neuropharm.2018.11.032.

Kovács, Richard, Zoltan Gerevich, Alon Friedman, Jakub Otáhal, Ofer Prager, Siegrun Gabriel, and Nikolaus Berndt. 2018. ‘Bioenergetic Mechanisms of Seizure Control’. Frontiers in Cellular Neuroscience 12 (October). 10.3389/fncel.2018.00335.

Kuramoto, Yoshiki. 1984. Chemical Oscilations Waves and Turbulence. Vol. 19. Springer Berlin Heidelberg.

Latham, P E, B J Richmond, P G Nelson, And S Nirenberg, and S Nirenberg. 2000. ‘Intrinsic Dynamics in Neuronal Networks. I. Theory’.

Levenstein, Daniel, György Buzsáki, and John Rinzel. 2019. ‘NREM Sleep in the Rodent Neocortex and Hippocampus Reflects Excitable Dynamics’. Nature Communications 10 (1). 10.1038/s41467-019-10327-5.

Linaro, Daniele, Marco Storace, and Maurizio Mattia. 2011. ‘Inferring Network Dynamics and Neuron Properties from Population Recordings’. Frontiers in Computational Neuroscience 5 (October). 10.3389/fncom.2011.00043.

Lőrincz, Magor L., David Gunner, Ying Bao, William M. Connelly, John T.R. Isaac, Stuart W. Hughes, and Vincenzo Crunelli. 2015. ‘A Distinct Class of Slow (∼0.2–2 Hz) Intrinsically Bursting Layer 5 Pyramidal Neurons Determines UP/DOWN State Dynamics in the Neocortex’. Journal of Neuroscience 35 (14): 5442–58. 10.1523/JNEUROSCI.3603-14.2015.

Marder, Eve, and Jean Marc Goaillard. 2006. ‘Variability, Compensation and Homeostasis in Neuron and Network Function’. Nature Reviews Neuroscience. 10.1038/nrn1949.

Massimini, Marcello, Maurizio Corbetta, Maria V. Sanchez-Vives, Thomas Andrillon, Gustavo Deco, Mario Rosanova, and Simone Sarasso. 2024. ‘Sleep-like Cortical Dynamics during Wakefulness and Their Network Effects Following Brain Injury’. Nature Communications 15 (1). 10.1038/s41467-024-51586-1.

Massimini, Marcello, Reto Huber, Fabio Ferrarelli, Sean Hill, and Giulio Tononi. 2004. ‘The Sleep Slow Oscillation as a Traveling Wave’. Journal of Neuroscience 24 (31): 6862–70. 10.1523/JNEUROSCI.1318-04.2004.

Mattia, Maurizio, Stefano Ferraina, and Paolo Del Giudice. 2010. ‘Dissociated Multi-Unit Activity and Local Field Potentials: A Theory Inspired Analysis of a Motor Decision Task’. NeuroImage 52 (3): 812–23. 10.1016/j.neuroimage.2010.01.063.

Mattia, Maurizio, and Maria V. Sanchez-Vives. 2012. ‘Exploring the Spectrum of Dynamical Regimes and Timescales in Spontaneous Cortical Activity’. Cognitive Neurodynamics 6 (3): 239–50. 10.1007/s11571-011-9179-4.

Mcguire, Barbara A, Charles D Gilbert, Patricia K Rivlin, and Torsten N Wiesel. 1991. ‘Targets of Horizontal Connections in Macaque Primary Visual Cortex’. The Journal of Comparative Neurology 305 (November):370–92.

Mehrotra, Dhruv, Daniel Levenstein, Adrian J Duszkiewicz, Sofia Skromne Carrasco, Sam A Booker, Angelika Kwiatkowska, and Adrien Peyrache. 2023. ‘Hyperpolarization-Activated Currents Drive Neuronal Activation Sequences in Sleep’. BioRxiv, 2023.09.12.557442. https://www.biorxiv.org/content/10.1101/2023.09.12.557442v1.

Mohajerani, Majid H., David A. McVea, Matthew Fingas, and Timothy H. Murphy. 2010. ‘Mirrored Bilateral Slow-Wave Cortical Activity within Local Circuits Revealed by Fast Bihemispheric Voltage-Sensitive Dye Imaging in Anesthetized and Awake Mice’. Journal of Neuroscience 30 (10): 3745–51. 10.1523/JNEUROSCI.6437-09.2010.

Monti, Jaime M., and Daniel Monti. 2007. ‘The Involvement of Dopamine in the Modulation of Sleep and Waking’. Sleep Medicine Reviews. 10.1016/j.smrv.2006.08.003.

Murphy, Michael, Brady A. Riedner, Reto Huber, Marcello Massimini, Fabio Ferrarelli, and Giulio Tononi. 2009. ‘Source Modeling Sleep Slow Waves’. Proceedings of the National Academy of Sciences of the United States of America 106 (5): 1608–13. 10.1073/pnas.0807933106.

Nádasdy, Zoltá, Hajime Hirase, Andrá Czurkó, Jozsef Csicsvari, and György Buzsáki. 1999. ‘Replay and Time Compression of Recurring Spike Sequences in the Hippocampus’. The Journal of Neuroscience 21 (19): 94979507.

Narikiyo, Kimiya, Rumiko Mizuguchi, Ayako Ajima, Momoko Shiozaki, Hiroki Hamanaka, Joshua P. Johansen, Kensaku Mori, and Yoshihiro Yoshihara. 2020. ‘The Claustrum Coordinates Cortical Slow-Wave Activity’. Nature Neuroscience. 10.1038/s41593-020-0625-7.

Nghiem, Trang, Anh E., Núria Tort-Colet, Tomasz Górski, Ulisse Ferrari, Shayan Moghimyfiroozabad, Jennifer S. Goldman, Bartosz Teleńczuk, et al. 2020. ‘Cholinergic Switch between Two Types of Slow Waves in Cerebral Cortex’. Cerebral Cortex 30 (6): 3451–66. 10.1093/cercor/bhz320.

Nir, Yuval, Richard J. Staba, Thomas Andrillon, Vladyslav V. Vyazovskiy, Chiara Cirelli, Itzhak Fried, and Giulio Tononi. 2011. ‘Regional Slow Waves and Spindles in Human Sleep’. Neuron 70 (1): 153–69. 10.1016/j.neuron.2011.02.043.

Parga, N, and Larry F Abbott. 2007. ‘Network Model of Spontaneous Activity Exhibiting Synchronous Transitions between up and down States’. Frontiers in Neuroscience 1 (1). doi:10.3389/neuro.01.1.1.004.2007.

Paxinos, George., and Keith B. J.. Franklin. 2001. The Mouse Brain in Stereotaxic Coordinates. Academic Press.

Pazienti, Antonio, Andrea Galluzzi, Miguel Dasilva, Maria V. Sanchez-Vives, and Maurizio Mattia. 2022. ‘Slow Waves Form Expanding, Memory-Rich Mesostates Steered by Local Excitability in Fading Anesthesia’. IScience 25 (3). 10.1016/j.isci.2022.103918.

Picciotto, Marina R., Michael J. Higley, and Yann S. Mineur. 2012. ‘Acetylcholine as a Neuromodulator: Cholinergic Signaling Shapes Nervous System Function and Behavior’. Neuron 76 (1): 116–29. 10.1016/j.neuron.2012.08.036.

Porta, Leonardo Dalla, Almudena Barbero-Castillo, Jose Manuel Sanchez-Sanchez, and Maria V. Sanchez-Vives. 2023. ‘M-Current Modulation of Cortical Slow Oscillations: Network Dynamics and Computational Modeling’. PLoS Computational Biology 19 (7): 1–17. 10.1371/journal.pcbi.1011246.

Potjans, Tobias C., and Markus Diesmann. 2014. ‘The Cell-Type Specific Cortical Microcircuit: Relating Structure and Activity in a Full-Scale Spiking Network Model’. Cerebral Cortex 24 (3): 785–806. 10.1093/cercor/bhs358.

Prinz, Astrid A., Dirk Bucher, and Eve Marder. 2004. ‘Similar Network Activity from Disparate Circuit Parameters’. Nature Neuroscience 7 (12): 1345–52. 10.1038/nn1352.

Reig, R., M. Mattia, A. Compte, C. Belmonte, and M. V. Sanchez-Vives. 2010. ‘Temperature Modulation of Slow and Fast Cortical Rhythms’. Journal of Neurophysiology 103 (3): 1253–61. 10.1152/jn.00890.2009.

Roš, Hana, Robert N.S. Sachdev, Yuguo Yu, Nenad Šestan, and David A. McCormick. 2009. ‘Neocortical Networks Entrain Neuronal Circuits in Cerebellar Cortex’. Journal of Neuroscience 29 (33): 10309–20. 10.1523/JNEUROSCI.2327-09.2009.

Rosanova, M., M. Fecchio, S. Casarotto, S. Sarasso, A. G. Casali, A. Pigorini, A. Comanducci, et al. 2018. ‘Sleep-like Cortical OFF-Periods Disrupt Causality and Complexity in the Brain of Unresponsive Wakefulness Syndrome Patients’. Nature Communications 9 (1): 1–10. 10.1038/s41467-018-06871-1.

Ruiz-Mejias, Marcel, Laura Ciria-suarez, Maurizio Mattia, Maria V Sanchez-vives, Marcel Ruiz-mejias, Laura Ciria-suarez, Maurizio Mattia, and Maria V Sanchez-vives. 2011. ‘Slow and Fast Rhythms Generated in the Cerebral Cortex of the Anesthetized Mouse’. J Neurophysiol 106 (August 2011): 2910–21. 10.1152/jn.00440.2011.

Russo, S., A. Pigorini, E. Mikulan, S. Sarasso, A. Rubino, F. M. Zauli, S. Parmigiani, et al. 2021. ‘Focal Lesions Induce Large-Scale Percolation of Sleep-like Intracerebral Activity in Awake Humans’. NeuroImage 234 (February): 117964. 10.1016/j.neuroimage.2021.117964.

Sanchez-Vives, Maria V., V. F. Descalzo, R. Reig, N. A. Figueroa, A. Compte, and R. Gallego. 2008. ‘Rhythmic Spontaneous Activity in the Piriform Cortex’. Cerebral Cortex 18 (5): 1179–92. 10.1093/cercor/bhm152.

Sanchez-Vives, Maria V., and M. Mattia. 2014. ‘Slow Wave Activity as the Default Mode of the Cerebral Cortex’. Archives Italiennes de Biologie 152 (2–3): 147–55. 10.12871/000298292014239.

Sanchez-Vives, Maria V., and David A. McCormick. 2000. ‘Cellular and Network Mechanisms of Rhytmic Recurrent Activity in Neocortex’. Nature Neuroscience 3 (10): 1027–34. 10.1038/79848.

Sanchez-Vives, Maria V, Ramon Reig, Milena Winograd, and Vanessa F Descalzo. 2007. ‘An Active Cortical Network in Vitro’. Research Signpost. Vol. 37.

Sanchez-Vives, Maria Victoria, Marcello Massimini, and Maurizio Mattia. 2017. ‘Shaping the Default Activity Pattern of the Cortical Network’. Neuron 94 (5): 993–1001. 10.1016/j.neuron.2017.05.015.

Sanchez-Vives, MV., M. Mattia, A. Compte, M. Perez-Zabalza, M. Winograd, V. F. Descalzo, and R. Reig. 2010. ‘Inhibitory Modulation of Cortical Up States’. Journal of Neurophysiology 104 (3): 1314–24. 10.1152/jn.00178.2010.

Sarasso, Simone, Sasha D’Ambrosio, Matteo Fecchio, Silvia Casarotto, Alessandro Viganò, Cristina Landi, Giulia Mattavelli, et al. 2020. ‘Local Sleep-like Cortical Reactivity in the Awake Brain after Focal Injury’. Brain 143 (12): 3672–84. 10.1093/brain/awaa338.

Schmidt, Stephen L., Erin Y. Chew, Davis V. Bennett, Mohamed A. Hammad, and Flavio Fröhlich. 2013. ‘Differential Effects of Cholinergic and Noradrenergic Neuromodulation on Spontaneous Cortical Network Dynamics’. Neuropharmacology 72:259–73. 10.1016/j.neuropharm.2013.04.045.

Schulz, David J., Jean Marc Goaillard, and Eve Marder. 2006. ‘Variable Channel Expression in Identified Single and Electrically Coupled Neurons in Different Animals’. Nature Neuroscience 9 (3): 356–62. 10.1038/nn1639.

Sheroziya, Maxim, and Igor Timofeev. 2014. ‘Global Intracellular Slow-Wave Dynamics of the Thalamocortical System’. Journal of Neuroscience 34 (26): 8875–93. 10.1523/JNEUROSCI.4460-13.2014.

Shimaoka, Daisuke, Chenchen Song, and Thomas Knöpfel. 2017. ‘State-Dependent Modulation of Slow Wave Motifs towards Awakening’. Frontiers in Cellular Neuroscience 11 (April). 10.3389/fncel.2017.00108.

Steriade, M. 2001. ‘Impact of Network Activities on Neuronal Properties in Corticothalamic Systems’. Journal of Neurophysiology 86 (1): 1–39. 10.1152/jn.2001.86.1.1.

Steriade, M, A Nuñez, and F Amzica. 1993. ‘A Novel Slow (< 1 Hz) Oscillation of Neocortical Neurons in Vivo: Depolarizing and Hyperpolarizing Components.’ The Journal of Neuroscience: The Official Journal of the Society for Neuroscience 13 (8): 3252–65. 3252-3265.

Stimberg, Marcel, Romain Brette, and Dan F.M. Goodman. 2019. ‘Brian 2, an Intuitive and Efficient Neural Simulator’. ELife 8:1–41. 10.7554/eLife.47314.

Strogatz, Steven H. 2000. ‘From Kuramoto to Crawford: Exploring the Onset of Synchronization in Populations of Coupled Oscillators’. Physica D. Vol. 143.

Stroh, Albrecht, Helmuth Adelsberger, Alexander Groh, Charlotta Rühlmann, Sebastian Fischer, Anja Schierloh, Karl Deisseroth, and Arthur Konnerth. 2013. ‘Making Waves: Initiation and Propagation of Corticothalamic Ca2+ Waves In Vivo’. Neuron 77 (6): 1136–50. 10.1016/j.neuron.2013.01.031.

Timofeev, I. 2000. ‘Origin of Slow Cortical Oscillations in Deafferented Cortical Slabs’. Cerebral Cortex 10 (12): 1185–99. 10.1093/cercor/10.12.1185.

Timofeev, I., and M. Steriade. 1996. ‘Low-Frequency Rhythms in the Thalamus of Intact-Cortex and Decorticated Cats’. Journal of Neurophysiology 76 (6): 4152–68. 10.1152/jn.1996.76.6.4152.

Tononi, Giulio, and Chiara Cirelli. 2006. ‘Sleep Function and Synaptic Homeostasis’. Sleep Medicine Reviews 10 (1): 49–62. 10.1016/j.smrv.2005.05.002.

Torao-Angosto, Melody, Arnau Manasanch, Maurizio Mattia, and Maria V. Sanchez-Vives. 2021. ‘Up and Down States During Slow Oscillations in Slow-Wave Sleep and Different Levels of Anesthesia’. Frontiers in Systems Neuroscience 15 (February): 1–9. 10.3389/fnsys.2021.609645.

Vinci, Gianni V., Roberto Benzi, and Maurizio Mattia. 2023. ‘Self-Consistent Stochastic Dynamics for Finite-Size Networks of Spiking Neurons’. Physical Review Letters 130 (9). 10.1103/PhysRevLett.130.097402.

Vyazovskiy, Vladyslav V., and Kenneth D. Harris. 2013. ‘Sleep and the Single Neuron: The Role of Global Slow Oscillations in Individual Cell Rest’. Nature Reviews Neuroscience 14 (6): 443–51. 10.1038/nrn3494.

Vyazovskiy, Vladyslav V., Umberto Olcese, Erin C. Hanlon, Yuval Nir, Chiara Cirelli, and Giulio Tononi. 2011. ‘Local Sleep in Awake Rats’. Nature 472 (7344): 443–47. 10.1038/nature10009.

Wang, Xiao Jing, and György Buzsáki. 1996. ‘Gamma Oscillation by Synaptic Inhibition in a Hippocampal Interneuronal Network Model’. Journal of Neuroscience 16 (20): 6402–13. 10.1523/jneurosci.16-20-06402.1996.

Wang, Yun, Henry Markram, Philip H. Goodman, Thomas K. Berger, Junying Ma, and Patricia S. Goldman-Rakic. 2006. ‘Heterogeneity in the Pyramidal Network of the Medial Prefrontal Cortex’. Nature Neuroscience 9 (4): 534–42. 10.1038/nn1670.

Wester, Jason C., and Diego Contreras. 2012. ‘Columnar Interactions Determine Horizontal Propagation of Recurrent Network Activity in Neocortex’. Journal of Neuroscience 32 (16): 5454–71. 10.1523/JNEUROSCI.5006-11.2012.

Wilson, Charles J., and Yasuo Kawaguchi. 1996. ‘The Origins of Two-State Spontaneous Membrane Potential Fluctuations of Neostriatal Spiny Neurons’. Journal of Neuroscience 16 (7): 2397–2410. 10.1523/jneurosci.16-07-02397.1996.

Winograd, Milena, Alain Destexhe, and Maria V. Sanchez-Vives. 2008. ‘Hyperpolarization-Activated Graded Persistent Activity in the Prefrontal Cortex’. Proceedings of the National Academy of Sciences of the United States of America 105 (20): 7298–7303. 10.1073/pnas.0800360105.

Yousheng, Shu, Andrea Hasenstaub, and David A McCormick. 2003. ‘Turning on and off Recurrent Cortical Activity’. Nature 423 (6937): 283–88. 10.1038/nature01614.

